# A Bayesian Noisy Logic Model for Inference of Transcription Factor Activity from Single Cell and Bulk Transcriptomic Data

**DOI:** 10.1101/2023.05.03.539308

**Authors:** Argenis Arriojas, Susan Patalano, Jill Macoska, Kourosh Zarringhalam

## Abstract

The advent of high-throughput sequencing has made it possible to measure the expression of genes at relatively low cost. However, direct measurement of regulatory mechanisms, such as Transcription Factor (TF) activity is still not readily feasible in a high-throughput manner. Consequently, there is a need for computational approaches that can reliably estimate regulator activity from observable gene expression data. In this work, we present a noisy Boolean logic Bayesian model for TF activity inference from differential gene expression data and causal graphs. Our approach provides a flexible framework to incorporate biologically motivated TF-gene regulation logic models. Using simulations and controlled over-expression experiments in cell cultures, we demonstrate that our method can accurately identify TF activity. Moreover, we apply our method to bulk and single cell transcriptomics measurements to investigate transcriptional regulation of fibroblast phenotypic plasticity. Finally, to facilitate usage, we provide user-friendly software packages and a web-interface to query TF activity from user input differential gene expression data: https://umbibio.math.umb.edu/nlbayes/.

**Author Summary:** NextGen RNA sequencing (RNA-Seq) has enabled simultaneous measurement of the expression level of all genes. Measurements can be done at the population level or single-cell resolution. However, direct measurement of regulatory mechanisms, such as Transcription Factor (TF) activity, is still not possible in a high-throughput manner. As such, there is a need for computational models to infer regulator activity from gene expression data. In this work, we introduce a Bayesian methodology that utilizes prior biological knowledge on bio-molecular interactions in conjunction with readily available gene expression measurements to estimate TF activity. The Bayesian model naturally incorporates biologically motivated combinatorial TF-gene interaction logic models and accounts for noise in gene expression data as well as prior knowledge. The method is accompanied by efficiently implemented R and Python software packages as well as a user-friendly web-based interface that allows users to upload their gene expression data and run queries on a TF-gene interaction network to identify and rank putative transcriptional regulators. This tool can be used for a wide range of applications, such as identification of TFs downstream of signaling events and environmental or molecular perturbations, the aberration in TF activity in diseases, and other studies with ‘case-control’ gene expression data.

## Introduction

Gene regulation plays an essential role in many cellular processes, including metabolism, signal transduction, development, and cell fate (Lelli, Slattery, & Mann, 2012; de Jong, 2002). At the transcriptional level, regulation of genes is orchestrated by the concerted action between Transcription Factors (TFs), histone modifiers, and distal *cis*-regulatory elements to finely tune and modulate expression of genes (Wilkinson, Nakauchi, & Gottgens, 2017). Sequence specific TFs, which have affinity for specific DNA sequences, may bind to *cis*-regulatory elements at the enhancer or promoter region of genes to either activate (upregulate) or repress (downregulate) the expression of genes. Aberration in TF activity and the dysregulation of target genes have been implicated in many pathological states and human disease (Lee & Young, 2013). Activity of TFs can be triggered downstream of signaling events, which in turn may be activated in response to environmental and molecular perturbations (Barolo & Posakony, 2002). Perturbations in TF activity often result in modulation of gene expression. The technological advancements in high-throughput sequencing have made it possible to measure expression of genes at relatively low cost. However, direct measurement of regulatory mechanisms, such as TF protein expression and functional activity in a high-throughput manner is still not readily available. Consequently, there is a need for computational approaches that can identify active regulatory mechanisms from observable gene expression data.

The scientific community has proposed several computational algorithms and biophysical models to study the impact of TF activity on gene expression. Some of these algorithms use statistical and probabilistic approaches to infer TF activity and dynamics directly from gene expression data (Bonneau, et al., 2006; Asif & Sanguinetti, 2011; Ocone & Sanguinetti, 2011; Bulashevska & Eils, 2005; Veber, Guziolowski, Le Borgne, Radulescu, & Siegel, 2008; Segal, et al., 2003), and more recently (Schacht, Oswald, Eils, Eichmüller, & König, 2014; Jiang, Freedman, Liu, & Liu, 2015; Fröhlich, 2015; Alvarez, et al., 2016; Yu, et al., 2017; Keenan, et al., 2019), while others rely on biophysical approaches to model expression of genes based on known TF-gene interactions (Honkela, et al., 2010; Djordjevic, Sengupta, & Shraiman, 2003). Boolean networks and probabilistic extensions have also been used to model gene regulation (Hashimoto, et al., 2004; Friedman, 2003; Bar-Joseph, et al., 2003; Segal, Taskar, Gasch, Friedman, & Koller, 2001; Segal, et al., 2003; Bulashevska & Eils, 2005). In logic models, genes are assumed to be either ON or OFF and Boolean logic (AND, OR, NOR, etc.) is utilized to model combinatorial regulation. For example, (Bulashevska & Eils, 2005) introduced a Bayesian approach to generalize the Boolean logic to incorporate noise and utilized their approach to reconstruct gene regulatory networks in yeast.

Another class of algorithms use prior biological knowledge on biomolecular interactions to link a differential gene expression (DGE) profile to upstream regulators (e.g., TFs) (Zarringhalam, Enayetallah, Gutteridge, Sidders, & Ziemek, 2013; Fakhry, et al., 2016; Subramanian, et al., 2005; Chindelevitch, et al., 2012; Chindelevitch, Loh, Enayetallah, Berger, & Ziemek, 2012; Kramer, Green, Pollard, & Tugendreich, 2014; Farahmand, O’Connor, Macoska, & Zarringhalam, 2019). The essential ingredients of these type of algorithms are (1) an input DGE profile, (ii) a network of biomolecular interactions or pre-defined gene sets, and (iii) an inference algorithm to query the network. The output is a set of candidate regulators, pathways, or biological processes with associated probabilities or significance p-values. The DGE profile as obtained from RNA-Seq or microarrays studies is the observable input that quantifies the difference in transcript abundance between two conditions (e.g., healthy vs. disease, stimulated vs. not stimulated, etc.). The network of biomolecular interactions encapsulates the prior biological knowledge. The inference algorithms typically map the DGE profile to the network to identify drivers (nodes, terms, or paths in the graph) of the observed transcriptional changes.

Despite the popularity and success of these methods, several challenges remain to be sufficiently addressed. For example, biophysical models are computationally expensive and are suitable for small scale applications or simulation studies. Boolean logic models, although simple to implement, cannot directly account for noise in gene expression data. On the other hand, probabilistic models for inference of gene regulatory networks typically overlook the context of experiment. Regulatory networks may be noisy or contain interactions that are applicable in a specific context only. Properly modeling any dependencies on the biological context in active regulator inference and enrichment analysis algorithms can lead to more accurate inference of the regulatory mechanisms specific to that context. Utilizing causality and information on mode of regulation (activation vs. repression) can also significantly reduce false positive predictions, resulting in more interpretable models. Moreover, biologically motivated TF-gene interaction logic models (e.g., combinatorial effects of activators and repressors on gene expression) must be taken into consideration when inferring transcriptional regulatory programs. To address these challenges, we have developed a Noisy-Logic Bayesian (NLBayes) TF activity inference model that accounts for these factors in a unified manner. Given an input DGE profile, our model incorporates the prior information on causal regulatory interactions and makes posterior adjustments to further account for noise and determine the context-specific posterior network structure and active regulators through a Gibbs sampling procedure.

We evaluate the performance of our model using simulation studies as well as over-expression datasets and demonstrate that our method can accurately identify active transcriptional regulators from gene expression data and causal graphs. We benchmark our algorithm against VIPER, a closely related method that is widely used for identification of regulon activity (Alvarez, et al., 2016). Both methods are able to identify relevant TFs in the corresponding experiments, with several TFs identified by both methods at the intersection, indicating that the algorithms complement each other. Our method can be used for novel biological discoveries. To illustrate this, we apply our method to investigate transcriptional regulation of fibroblast phenotypic plasticity in response to signaling molecules TGFβ and CXCL12. This study utilizes differential expression profiles from bulk RNA-Seq experiment using prostate cell lines (Gharaee-Kermani, et al., 2012), stimulated with TGFβ and CXCL12. Our analysis recovers several TFs, including YAP1 as the top prediction, which has been identified as a driver of myofibroblast differentiation in multiple tissue phenotypes (Piersma, et al., 2015; Pang, et al., 2023; Li, et al., 2022; Li, et al., 2022; Xu, et al., 2021; Lee, et al., 2022; Salloum, et al., 2021; Wang, et al., 2020; Li, et al., 2021; Allison, 2021). Additionally, we present new single cell gene expression data from the same prostate cell lines as well as three additional human prostate fibroblast cell lines. We characterize the cell lines at the transcriptional level and apply our algorithm to identify major transcriptional regulators in each cell line and study the impact of immortalization on transcriptional regulation. Our algorithm provides a general framework and a widely applicable tool to study transcriptional regulators of differential gene expression. To facilitate wider use, we provide R and python packages, and a web-interface for running inference experiments. **Fig. 1** summarizes the overall approach.

**Fig. 1.**
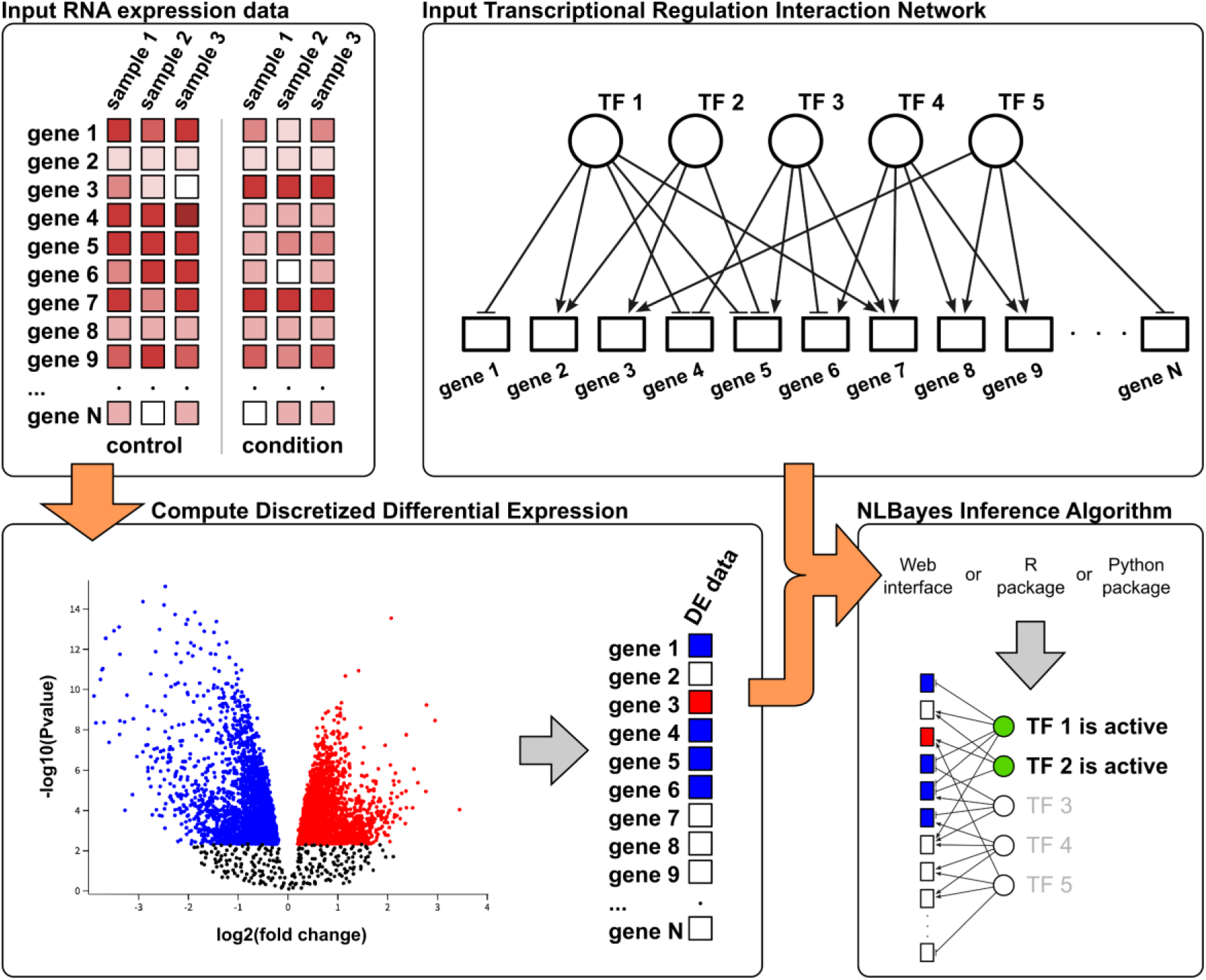
Schematic diagram of the NLBayes inference pipeline. The starting point is gene expression data from two conditions, from which a differential gene expression profile is calculated by discretizing gene values to -1 (down regulated), 0 (not regulated) or 1 (up regulated) using cutoff thresholds on p-values and/or foldchange. A TF-gene interaction network is used to build the graphical model. Values for gene nodes are populated from the differential expression profile. NLBayes runs a probabilistic query on the causal network and outputs the posterior distribution of TFs, from which the activation state of the TFs is determined.

## Results

To test our approach, we performed a series of experiments including simulations studies, benchmarks against an alternative approach, and inference of TF activity on novel datasets.

The core of the algorithm uses an OR-NOR transcriptional regulation logic to predict TF activity. In this logic two conditions must be satisfied for gene activation: 1) at least one of activator is targeting the gene and 2) no inhibitor is targeting the gene. On the other hand, for down regulation of genes we use a simple OR transcriptional regulation logic model: at least one inhibitor must target the genes.

The algorithm outputs posterior probabilities for each TF activation state. The prior probability for TF activation is set to a small value (*p*_0_ = 0.01), and as such we consider a TF with posterior probability *p* ≥ 0.2 as potentially relevant. We then define three thresholds to classify inferred active TFs: High-confidence (*p* ≥ 0.8) Mid-confidence (*p* ≥ 0.5), and Low-confidence (*p* ≥ 0.2).

### Simulation studies

We performed several simulation studies to assess the ability of our algorithm in recovering active transcriptional regulators from gene expression data and to test the robustness of the inference process to noise in gene expression data and the causal graph of TF-gene interactions. For this analysis we generated a random interaction network consisting of 250 TFs and 5000 downstream target genes. The downstream targets of each TF were picked at random from a binomial distribution, resulting in ∼30000 interactions (edges) in the network. The edges of the network were randomly set as activation (65%) and inhibition (35%).

We randomly selected 10 TFs and assigned them as active (the ground truth) and simulated downstream differential gene expression data by assigning +1 or -1 to 10% of the target genes according to the causal graph. For genes targeted by multiple active TFs, we calculated the algebraic sum of all incoming interactions and took the net sign. This produced an average of 120 differentially expressed genes. Each experiment was repeated 20 times.

#### Impact of data randomization

For this analysis, we randomized a fraction of the input data (0, 0.25, 0.5, 0.75, 1.00) by randomly toggling the values. At 0% the data is not randomized, while at 100% the input data is completely random. The inference procedure was run on each input data set. **Figs. 2A and 2B** show the ROC and precision-recall curves for each randomization experiment respectively. **Fig. 2C** illustrates the sampled posterior distributions for active TFs at different randomization levels. As expected, at 100% the model fails to recover active TFs. However, up to 75% randomization the algorithm is still able to recover active TFs, with significant increase in accuracy at lower levels of noise.

**Fig. 2.**
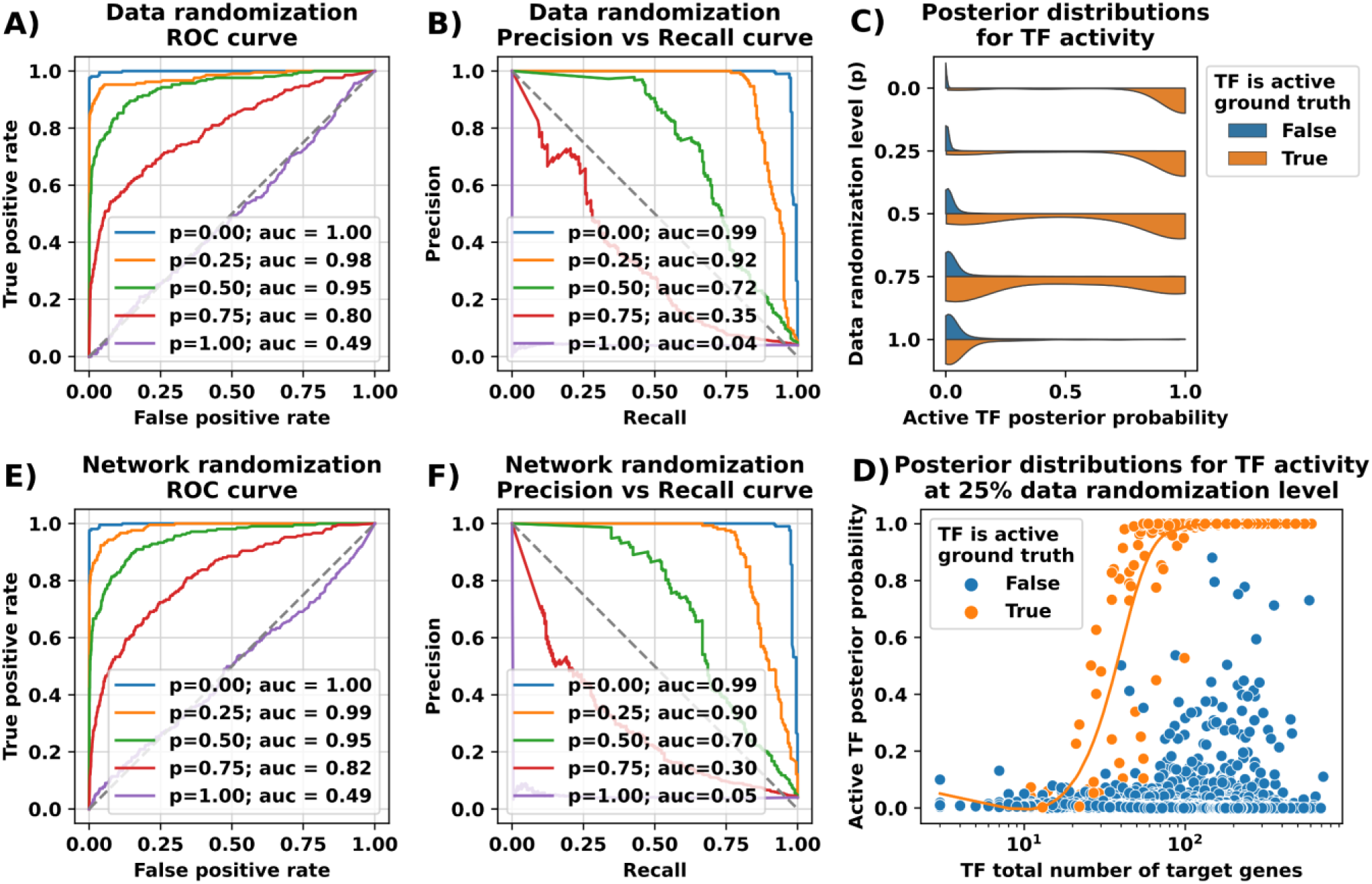
Performance evaluation of the model before and after randomization of data. ROC curves and Precision vs Recall curves are shown for randomization simulation in input gene expression (A, B) and TF-gene interaction network (E, F). p indicates the corresponding fraction of randomization used. AUC scores are displayed in the legend. (C) The posterior distributions for TF activity is shown. The colors indicate the ground truth, i.e. whether the TF was set to active in the simulation. (F) The impact of the number of target genes on the inference results for the case of 25% randomization of data.

To assess the impact of the number of target genes regulated by a TF on the posterior probability of the TF’s activity, we ran multiple simulations and plotted the posterior probability vs the number of genes regulated by TFs, color coded by TF activity (**Fig. 2D).** For these simulations, we used noisy data at 25% randomization. We observe that for active TFs with less than 30 target genes, the posterior probability is low. This is expected as in these simulations, only 10% of target genes are set as differentially expressed, yielding an average of only 3 target genes that are modulated. This information is too small to shift the posterior probability. However, as the total number of target genes increase, we observe a threshold effect, where posterior probabilities stabilize, and the total number of targets does not have an impact on the inference. These experiments were repeated for several other randomly generated networks with consistent results (SI File Table S4).

#### Impact of network randomization

For this analysis, we select a fraction (0, 0.25, 0.5, 0.75, 1.00) of the edges in the network and randomly reassign them to different target genes. The inference was run using unperturbed input gene expression data and randomized networks. **Figs. 2E and 2F** show the ROC and precision-recall curves for each randomization experiment. A similar picture as in data randomization emerges.

These results demonstrate the robustness of our algorithm at 25% randomization level in both the gene expression data as well as the input network, while still retaining some prediction power at the 50% and 75% randomization levels.

### Over-expression datasets

To test the ability of our algorithm in recovering known active regulators, we used three publicly available over-expression datasets (GSE3151), performed on human primary mammary epithelial cell cultures, each generated by over-expression of an oncogene: E2F3, c-Myc, and H-Ras (Bild, et al., 2006). For these inference experiments, we utilized a TF-gene interaction network generated by Farahmand et. al. (Farahmand, O’Connor, Macoska, & Zarringhalam, 2019). The network was generated using several high-throughput datasets, and a Gaussian Graphical Model. We chose to use this network as it showed consistently high predictive power across several datasets (Farahmand, O’Connor, Macoska, & Zarringhalam, 2019). In each experiment, differentially expressed genes were determined compared to the control sample and inputted into the TF activity inference algorithm. **Table 1** summarizes the inference results. Inferred active regulators are split into three categories based on posterior probability *p* of inferred activity: High-confidence (*p* ≥ 0.8) Mid-confidence (*p* ≥ 0.5), and Low-confidence (*p* ≥ 0.2). Percentage of differentially expressed genes targeted (explained) by at least one inferred active regulator is also presented in the table.

**Table 1.** Predicted active TFs from 3 over-expression experiments: E2F, MYC, and RAS. Top 3 rows summarize the total number of Differentially Expressed Genes (DEGs) and the number of genes with increased and decreased RNA expression. Rows 4 and 5 show the proportion of explained DEGs by at least 1 inferred regulator at the indicated confidence level. Bottom colored panel lists the inferred regulators, split into three categories, high-confidence (green), mid-confidence (blue) and low-confidence (yellow).

| E2F3 |  |  | MYC |  |  | RAS |  |  |
| --- | --- | --- | --- | --- | --- | --- | --- | --- |
| max p-value = 0.01<br>min abs(log2 fold-change) = 1.00 |  |  | max p-value = 0.01<br>min abs(log2 fold-change) = 1.00 |  |  | max p-value = 0.01<br>min abs(log2 fold-change) = 2.00 |  |  |
| DEG | Increased | Decreased | DEG | Increased | Decreased | DEG | Increased | Decreased |
| 418 | 321 | 97 | 127 | 75 | 52 | 409 | 163 | 246 |
| Confidence level |  | DEG explained | Confidence level |  | DEG explained | Confidence level |  | DEG explained |
| 0.80 |  | 53% | 0.80 |  | 32% | 0.80 |  | 58% |
| 0.50 |  | 61% | 0.50 |  | 49% | 0.50 |  | 64% |
| 0.20 |  | 65% | 0.20 |  | 70% | 0.20 |  | 71% |
| Name | Enrichment | Inference | Name | Enrichment | Inference | Name | Enrichment | Inference |
| E2F1 | 3.7E-06 | 1.00 | TCF3 | 4.9E-01 | 1.00 | E2F3 | 6.1E-01 | 1.00 |
| FLI1 | 2.7E-02 | 1.00 | HDAC2 | 1.0E+00 | 0.90 | PPARG | 4.0E-01 | 1.00 |
| FOXA2 | 7.0E-01 | 1.00 | TP53 | 1.2E-02 | 0.85 | TEAD4 | 4.9E-01 | 1.00 |
| HOXA4 | 7.0E-01 | 1.00 | - | - | - | KLF5 | 2.2E-02 | 1.00 |
| MXI1 | 2.6E-01 | 1.00 | - | - | - | FOSL1 | 2.4E-06 | 1.00 |
| BCOR | 2.2E-02 | 1.00 | - | - | - | RELA | 1.4E-01 | 1.00 |
| AR | 1.4E-02 | 1.00 | - | - | - | ASCL1 | 1.0E+00 | 0.99 |
| FOXF1 | 7.9E-01 | 1.00 | - | - | - | STAT3 | 6.1E-01 | 0.99 |
| TCF3 | 8.6E-01 | 1.00 | - | - | - | PRKDC | 1.0E+00 | 0.97 |
| MBD3 | 1.0E+00 | 0.99 | - | - | - | JUND | 1.4E-01 | 0.97 |
| SOX2 | 7.3E-01 | 0.97 | - | - | - | NFKB2 | 1.2E-02 | 0.96 |
| - | - | - | - | - | - | SOX2 | 5.3E-01 | 0.96 |
| - | - | - | - | - | - | CTCFL | 1.0E+00 | 0.94 |
| - | - | - | - | - | - | GATA3 | 9.0E-01 | 0.92 |
| - | - | - | - | - | - | KDM6B | 3.2E-03 | 0.86 |
| HNF1B | 1.0E+00 | 0.77 | SIX2 | 1.0E+00 | 0.77 | MYC | 2.1E-01 | 0.75 |
| FOXM1 | 1.7E-02 | 0.76 | ETV1 | 1.0E+00 | 0.73 | NME2 | 1.0E+00 | 0.66 |
| NR5A2 | 9.2E-01 | 0.72 | KLF11 | 1.0E+00 | 0.54 | ESRRA | 1.0E+00 | 0.65 |
| HNF4A | 1.0E+00 | 0.71 | ILF3 | 2.9E-01 | 0.51 | SREBF1 | 1.8E-01 | 0.64 |
| POU5F1 | 7.0E-01 | 0.69 | - | - | - | MBD2 | 9.7E-01 | 0.64 |
| GATA3 | 1.5E-01 | 0.68 | - | - | - | - | - | - |
| TCF21 | 1.0E+00 | 0.50 | - | - | - | - | - | - |
| PGR | 8.6E-01 | 0.39 | PAX5 | 1.0E+00 | 0.49 | PBX4 | 1.0E+00 | 0.41 |
| TEAD1 | 1.0E+00 | 0.39 | TFAP4 | 2.0E-01 | 0.49 | HOXB13 | 1.0E+00 | 0.37 |
| GATA6 | 1.0E+00 | 0.39 | ZC3H8 | 5.0E-01 | 0.45 | FOXF1 | 3.6E-03 | 0.36 |
| VEZF1 | 1.0E+00 | 0.30 | PPARG | 7.6E-01 | 0.44 | KLF6 | 1.8E-02 | 0.34 |
| EGR1 | 7.0E-01 | 0.29 | PDX1 | 6.9E-01 | 0.42 | BCOR | 9.7E-01 | 0.33 |
| NKX2-1 | 1.0E+00 | 0.29 | TEAD4 | 2.9E-01 | 0.37 | HNF1B | 1.0E+00 | 0.32 |
| HOXA9 | 4.3E-01 | 0.26 | BATF | 7.3E-01 | 0.37 | SRF | 9.0E-01 | 0.31 |
| HOXC5 | 1.0E+00 | 0.26 | THAP11 | 1.0E+00 | 0.36 | GATA4 | 1.0E+00 | 0.27 |
| - | - | - | SMARCC2 | 1.0E+00 | 0.33 | CEBPB | 4.1E-02 | 0.24 |
| - | - | - | NCOA1 | 1.0E+00 | 0.33 | MAFF | 3.6E-03 | 0.22 |
| - | - | - | TRIM28 | 2.9E-01 | 0.24 | - | - | - |
| - | - | - | MYB | 1.0E+00 | 0.21 | - | - | - |

For the E2F3 expression data, the E2F1 is returned as the top regulator. E2F1 and E2F3 have a similar function in control of the cell cycle and are similarly implicated in cancer (Chen, Tsai, & Leone, 2009). E2F3 also regulates expression of FLI1, an ETS domain transcription factor and proto-oncogene (Li, Luo, Liu, Zacksenhaus, & Ben-David, 2015), as well as the FOXA2 transcription factor that promotes aggressive prostate cancer (Qi, Pellecchia, & Ronai, 2010) and HOXA4, a transcription factor important for embryonic development but often over-expressed in human colorectal (Bhatlekar, et al., 2014) and ovarian (Yamashita, et al., 2006) cancers.

The MYC and RAS oncogenic proteins transcriptionally activate multiple genes associated with tumor progression and prognosis. MYC transcriptionally activates TCF3, HDAC2 and TP53. The TCF3 transcription factor was recently found to promote gastric (Xie, et al., 2021) and endometrial (Gui, et al., 2021) cancers, among others. HDAC2 has been reported to promote metastasis in pancreatic (Krauß, et al., 2021) and breast (Huang, et al., 2021) cancers. MYC also regulates transcription of the well-known TP53 tumor suppressor gene. RAS activates transcription of E2F3 itself, as well as PPARG and TEAD4. Modulation of PPARG activity has been intensively examined as an anti-cancer therapeutic target (Chi, et al., 2021), and TEAD4 which is known to modulate different cellular processes in cancer via its transcriptional output (Chen, et al., 2020).

Taken together, these results demonstrate that the algorithm can accurately detect modulated transcriptional signals from DNA binding proteins.

### Benchmarks

We compared the performance of our algorithm against VIPER (Alvarez, et al., 2016), a widely used method for inference of regulon activity input gene expression data and causal graphs. For this benchmark, we used the Human Breast Carcinoma context specific network from ARACNe interactome (Lachmann, Giorgi, Lopez, & Califano, 2016). This network is appropriate for overexpression datasets as they used human breast cell cultures and allows a fair comparison. **Fig. 3** summarizes the overlap between the two algorithms. Overall, there is good agreement between the methods as well as regulators recovered only by one algorithm, demonstrating the viability of both methods in recovering modulated TFs. Each algorithm predicts TF activity that is not shared by the other algorithm. In the E2F3 overexpression experiment, our algorithm infers SOX17 as an active TF. SOX genes have been shown to interact with E2F3, and to be regulated by common micro-RNAs. For example, MicroRNA-141 regulates both SOX17 and E2F3 (Hamidi, Taghehchian, Basirat, Zangouei, & Moghbeli, 2022; Zhou, et al., 2015; Jia, et al., 2012). On the other hand, VIPER detects MEIS2 as a relevant TF in the overexpression of E2F3. siRNA-mediated silencing of RB1 and MEIS2 increases expression of E2F3 (Alam, et al., 2019). This shows that both approaches can complement each other, providing a wider picture of TF activity. While individually predicted TFs by each method may be important and can provide useful clues to the underlying biology, there is higher confidence in predictions at the intersection of both methods.

**Fig. 3.**
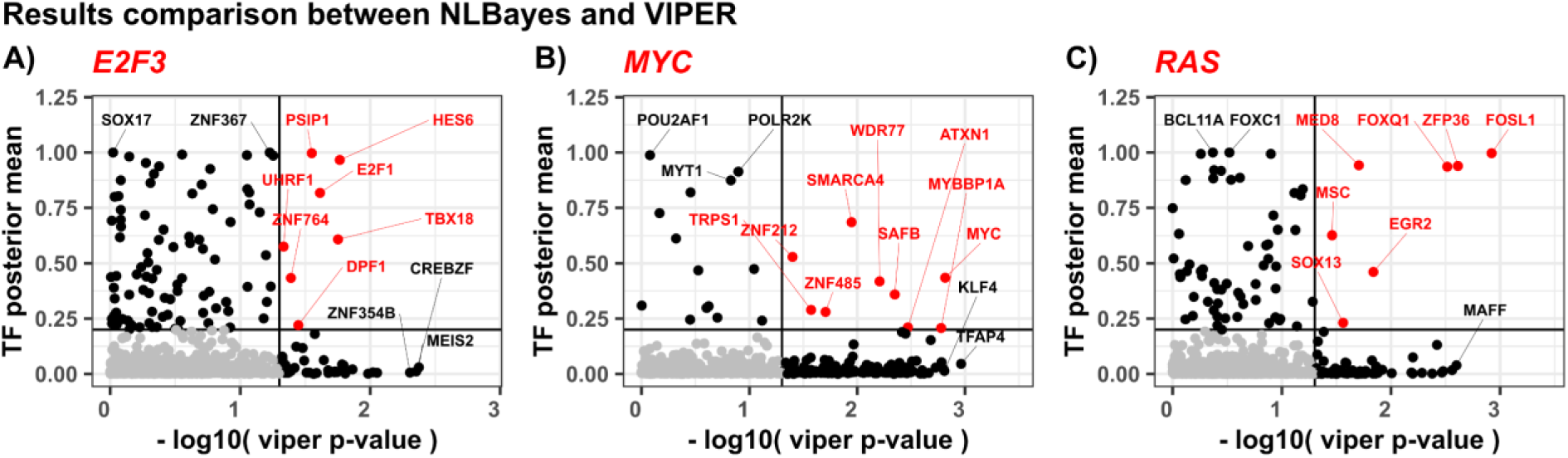
Comparison of active TF predictions by our method (y axis) and VIPER (x axis), in three separate overexpression experiments. Input network used is the BRCA derived regulon from (Alvarez, et al., 2016). Jointly predicted regulators are colored in red. Top predictions specific to one algorithm are labeled in black. Gray area shows low confidence predictions by both algorithms.

### Fibroblast phenotypic plasticity

To demonstrate the utility of our methodology in discovery of novel biology, we applied our algorithm to study fibroblast-to-myofibroblast phenotypic conversion in response to pro-fibrotic signaling molecules TGFβ and CXCL12 (Patalano, et al., 2018; Gharaee-Kermani, et al., 2012; Rodríguez-Nieves, Patalano, Almanza, Gharaee-Kermani, & Macoska, 2016). In this experiment, patient derived, immortalized prostate N1 cells were treated with the pro-fibrotic proteins TGFβ and CXCL12, both of which are known to promote collagen expression (Patalano, et al., 2018). TGFβ acts upon TGF/TGFR signaling axis and activates multiple Smad proteins, while CXCL12 acts upon CXCL12/CXCR4-axis, which transactivates EGFR and downstream signaling through MEK/ERK and PI3K/Akt pathways. Both signaling axes converge in the nucleus and promote the expression of multiple collagen genes (Gharaee-Kermani, et al., 2012; Rodríguez-Nieves, Patalano, Almanza, Gharaee-Kermani, & Macoska, 2016). RNA-Seq data was processed and compared to the background model to generate differential gene expression profiles as previously described (Patalano, et al., 2018). Differential gene expression data from TGFβ and CXCL12 treated cells were identified at fold change ≥ 2 and four cutoff thresholds for p-value. The different cutoffs for p-value were applied to examine the impact of stringency in significance on the inference results, and were chosen such that only 200, 400, 600 or 800 top differentially expressed genes are considered. To achieve this, we sorted the table of differentially expressed genes by p-values in ascending order and took the top rows for the analysis. Both datasets were used as input to the TF activity inference algorithm. For these experiments (and the remaining experiments), we used the 3-tissue TF-gene interaction network generated by Farahmand et. al. This network contains interaction edges that are common in at least three of the tissues used in that work and showed consistent performance across multiple datasets (Farahmand, O’Connor, Macoska, & Zarringhalam, 2019). **Table 2** summarizes the results. Inferred active regulators upon CXCL12 induction are largely similar to that of TGFβ (**Table 2**). This is expected as transcriptional profiles induced by TGFβ and CXCL12 are 75% similar (Patalano, et al., 2018). The top predicted regulators for TGFβ and CXCL12 for the top 200 DEGs are YAP1, RBPJ, KMT2C, ELF1, STAT1, and BPTF. YAP1 is known to play a role in the development and progression of multiple cancers as a transcriptional regulator of this signaling pathway and may function as a potential target for cancer treatment (Astudillo, 2022). Moreover, YAP1 has been identified as a driver of myofibroblast differentiation in several tissue phenotypes, like skin, heart, lung, pharynx, liver and kidney (Piersma, et al., 2015; Pang, et al., 2023; Li, et al., 2022; Li, et al., 2022; Xu, et al., 2021; Lee, et al., 2022; Salloum, et al., 2021; Wang, et al., 2020; Li, et al., 2021; Allison, 2021). STAT1 is a member of the STAT protein family. In response to cytokines and growth factors, STAT family members are phosphorylated by the receptor associated kinases, and then form homo- or heterodimers that translocate to the cell nucleus where they act as transcription activators. The protein encoded by this gene can be activated by various ligands including interferon-alpha, interferon-gamma, EGF, PDGF and IL6. This protein mediates the expression of a variety of genes, which is thought to be important for cell viability in response to different cell stimuli (Harrison & Moseley, 2020). RB1 is a negative regulator of the cell cycle and was the first tumor suppressor gene found. The encoded protein also stabilizes constitutive heterochromatin to maintain the overall chromatin structure. The active, hypophosphorylated form of the protein binds transcription factor E2F1. The deletion or mutation of the RB1 gene in many human cancers defines RB1 as a tumor suppressor gene (Knudsen, Pruitt, Hershberger, Witkiewicz, & Goodrich, 2019). LIN9 is a tumor suppressor protein that inhibits DNA synthesis and oncogenic transformation through association with the retinoblastoma 1 protein. The encoded protein also interacts with a complex of other cell cycle regulators to repress cell cycle-dependent gene expression in non-dividing cells (Walston, Iness, & Litovchick, 2021).

**Table 2.**
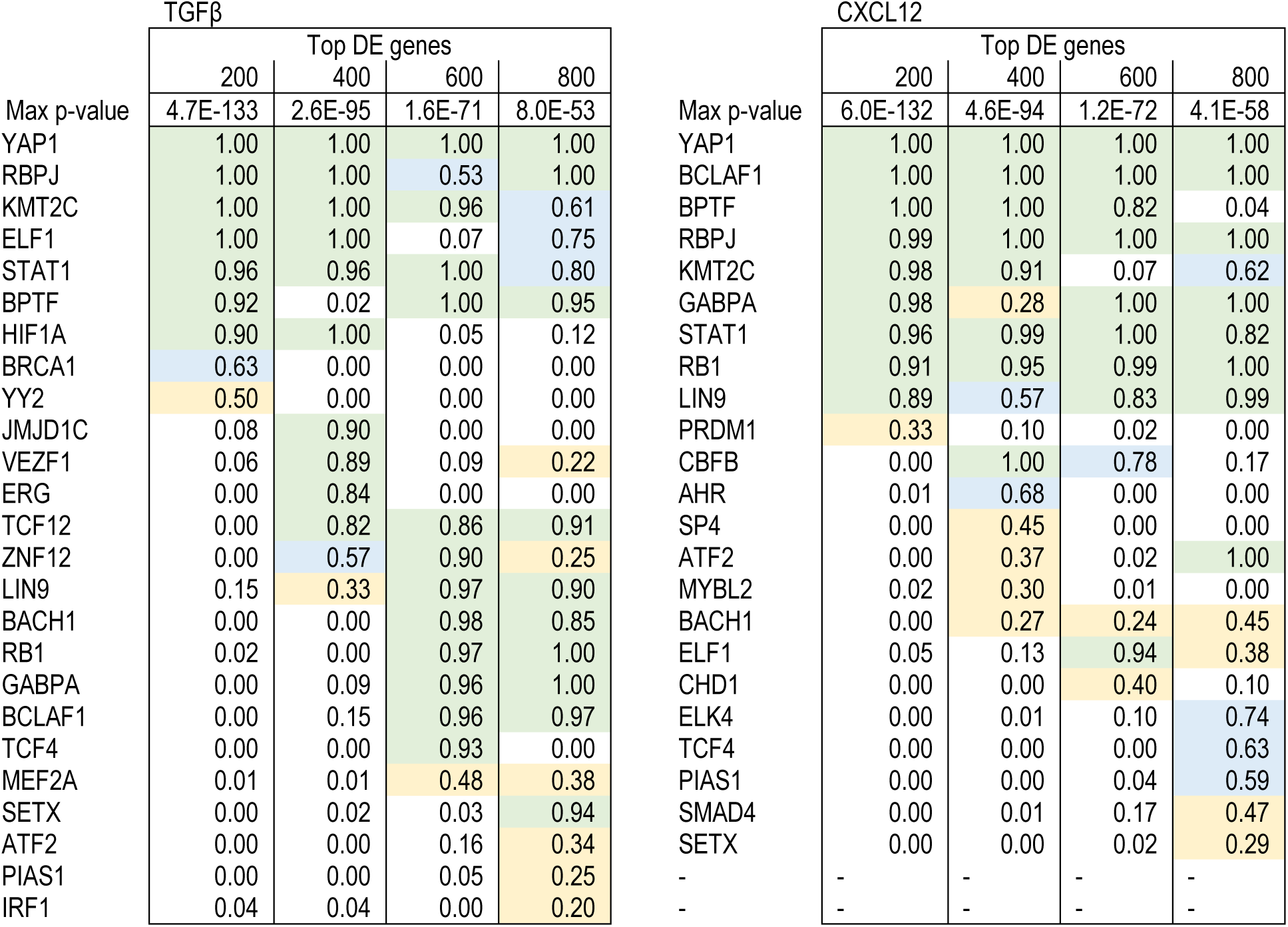
Predicted active TFs upon TGFβ (left) and CXCL12 induction (right). The top row shows total number of DEGs using four different p-value cutoff thresholds (Max p-value). Bottom colored panel lists the inferred regulators, split into three categories, high-confidence (green), mid-confidence (blue) and low-confidence (yellow).

We note that all these predicted active TFs appear as top regulators for the top 400, 600 and 800 DEGs, indicating the robustness of the algorithm to cutoff stringency criteria in input DGE profiles.

### Fibroblast heterogeneity: Single cell experiments

We performed several scRNA-seq experiments to further investigate the phenotypic plasticity of human prostate fibroblasts and characterize heterogeneity in cell populations. For this study, we utilized 4 human prostate cell lines: N1, SFT1, pHPF, and iHPF (For more information, see SI File Table S1). N1 cells are HPV E6/E7-immortalized prostate stromal fibroblasts originally explanted and grown from a stromal <u>N</u>odule of benign prostatic hyperplasia (Begley, Kasina, MacDonald, & Macoska, 2008). They exhibit a fibroblastic morphology, and express fibroblastic markers vimentin and calponin. These cells demonstrate secretion and proliferation profiles consistent with aging primary prostate fibroblasts. SFT1 cells are spontaneously immortalized prostate fibroblasts grown from a prostate of a patient with a <u>S</u>olitary <u>F</u>ibrous <u>T</u>umor of the prostate (Gharaee-Kermani, Mehra, Robinson, Wei, & Macoska, 2014). These cells carry an uncommon NAB2/STAT6 fusion gene that is associated with solitary fibrous tumors and likely accounted for cellular immortalization. pHPF cells are primary <u>H</u>uman <u>P</u>rostate <u>F</u>ibroblasts, purchased at passage 3 from Lifeline Cell Technology, harvested from young adult male. Finally, iHPF cells are created through transduction from pHPF cells with an EF1α-driven hTERT Lentivirus construct and have grown continuously in culture >30 passages.

We applied scRNA-seq to all 4 cell lines. **Fig. 4A** shows the UMAP projection of the cell lines. The N1 and SFT1 form distinct clusters in close proximity. Most of the cells are in G1 phase. pHPF cells also form a single cluster, mostly consisting of cells in G1 phase. Interestingly the iHPF cells cluster in two groups (A and B) that surround the primary pHPF cells. The majority of cells in iHPF_A cluster are in G1 phase, while iHPF_B consists of a mix of cells in G1, G2, S, and M phases. To further investigate the identity of these cells, we merged the data with FACS sorted single cell expression data derived from prostate tissue generated by Henry et. al. (Henry, et al., 2018). **Fig. 4B** shows that the RNA expression profiles of the five human prostate fibroblast cell lines N1, SFT1, iHPF_A and iHPF_B, and primary pHPF cells, cluster as expected with that of tissue-derived human prostate fibroblasts. As seen in **Fig. 4C**, the five human prostate fibroblast cell lines share a large signature of highly and commonly expressed genes, likely reflecting their common fibroblastic cell type. In particular, all five cell lines express COL1A1 (collagen 1) and VIM (vimentin). Examination of the top 10 differentially expressed genes in the four immortalized cell lines compared to primary human fibroblasts shows that N1 and SFT1 demonstrate a high degree of overlap and commonly express several inflammation-associated genes (CXCL1, ZNFAS1, CHI3L1). The iHPF_A and iHPF_B share a common gene signature as well that includes gene encoding signaling proteins (BEX1, WNT5A), growth factors and pathways (EREG, IGFBP5), and a gene over-expressed in the autoimmune disease, rheumatoid arthritis (TGM2). However, iHPF_B cells also highly express genes that are not expressed by iHPF_A, including several associated with vasculogenesis or angiogenesis (ANGPT1, F3, ADAMT1) or connective tissue and bone growth (TNFRSF11B). This suggests that iHPF_B cells may phenotypically resemble endothelial cells, which can differentiate from fibroblasts (Junker, et al., 2013). **Fig. 4E** quantifies the average log FC of top expressed markers (**Fig. 4C**) compared to the background (pHPF). Taken together, these data suggest that a seemingly homogenous culture of primary stromal prostate fibroblasts may comprise several subpopulations as have recently been shown for dermal fibroblasts (Hu, Moore, & Longaker, 2018; Philippeos, et al., 2018).

**Fig. 4.**
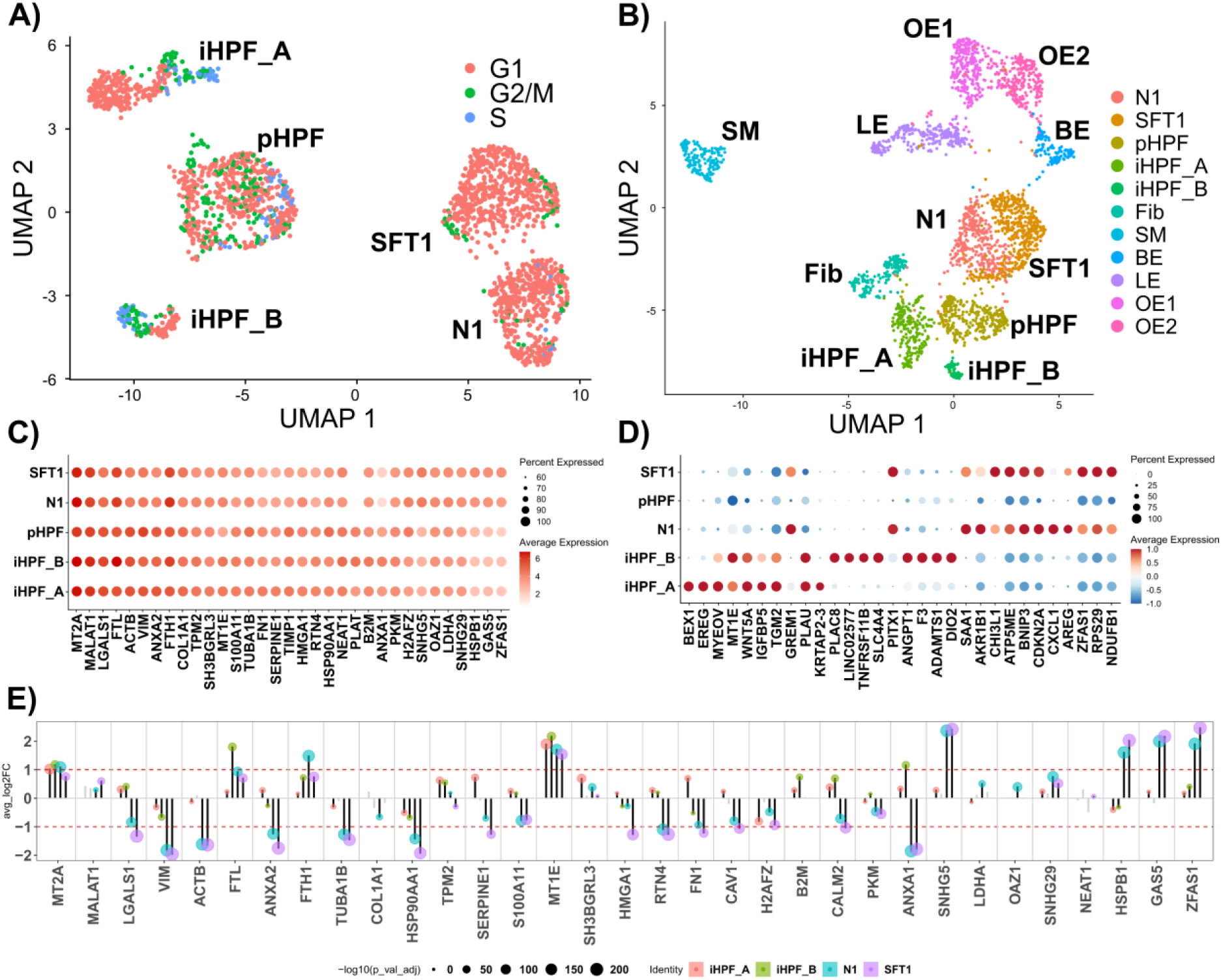
Single cell gene expression data from 4 prostate cell lines. A) UMAP projection of cell lines. iHPF shows two separate phenotypes here termed iHPF_A and iHPF_B. Cell cycle phase G1 is prominent in N1 and SFT1, but pHPF cells show significant number of cells in S and G2/M phases. B) Integration single prostate tissue data (Henry, et al., 2018), showing clusters for Fibroblasts (Fib), Smooth Muscle (SM), Basal Epithelia (BE), Luminal Epithelia (LE) and Other Epithelia (OE1, OE2). All 4 cell lines, N1, SFT1, pHPF and iHPF, appear as interconnected clusters, lying between epithelial and fibroblast cells. C) Top expressed genes. Overall, all 5 cell classes share the same highly expressed genes including COL1A1. D) Top 10 differentially expressed genes with respect to pHPF as a reference. Smallest and largest p-values are 1E-128 and 1E-26 respectively. See SI File Tables S2 and S3 for a list of top genes for each cell line, as shown in (C) and (D); Full list of differentially expressed genes is available in SI Table 1. E) Differential expression with respect to pHPF, for the highly expressed genes shown in C. FC of genes in panel C compared to pHPF. Notably, even though collagen expression high in across all groups, it is downregulated in N1 when compared to pHPF.

Next, we sought to quantitatively characterize similarities between cell lines at the transcriptional level. We first performed a differential gene expression analysis using the pHPF cell line as the background. **Fig. 5A** shows a bar plot of total number of upregulated and downregulated genes in each cell line compared to pHPF cells. We performed a gene set enrichment analysis on up & down regulated genes (**Fig. 5B**). As expected, the N1 and SFT1 cells demonstrate a high level of similarity, as we have previously shown that they respond similarly to stimulation with pro-fibrotics (Rodríguez-Nieves, Patalano, Almanza, Gharaee-Kermani, & Macoska, 2016). Conversely, although immortalized from the pHPF cells, iHPF_A and iHPF_B demonstrate a higher than expected dissimilarity, potentially reflecting fibroblast heterogeneity in the primary cell culture from which they were derived.

**Fig. 5.**
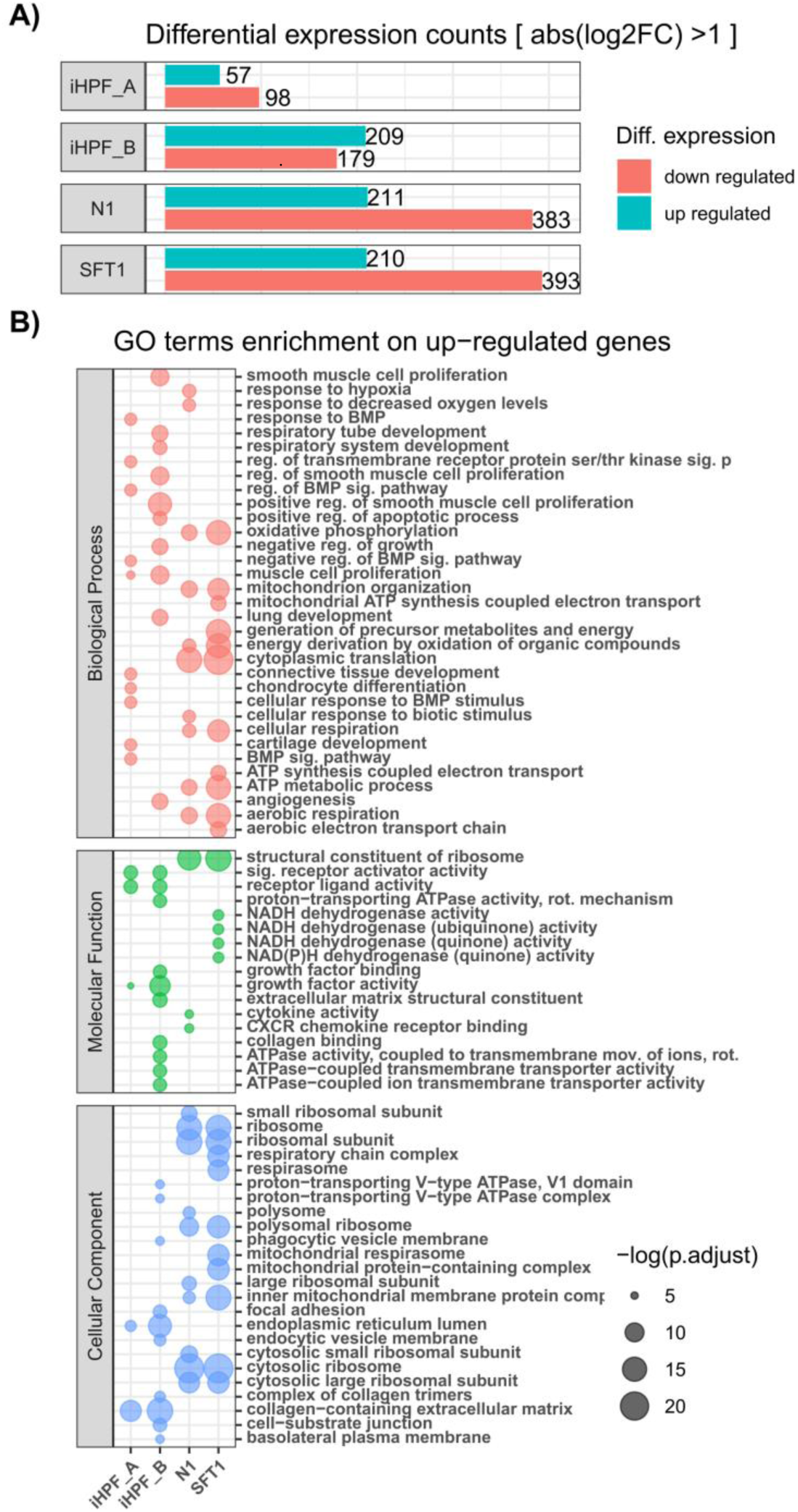
A) Total number of DEGs compared to the background model (pHPF). B) GO term Enrichment analysis of up regulated genes in each cell line (columns). See SI File Fig S1 for the GO term Enrichment analysis on down regulated genes.

Next, we applied our algorithm to the DEG profiles from each cell line to quantify similarity in transcriptional gene regulation. **Fig. 6** shows top inferred regulators in each cell line (left panel bar plots), along with their RNA expression level (middle panel bar plots), and the corresponding enrichment of the differentially expressed target genes (right panel bar plots). The enrichment analysis was performed by quantifying the overlap between the targets of the TF and DEGs using Fisher’s exact test. This analysis was performed for comparison of enrichment-based methods with our approach. Enrichment-based approaches do not consider the global topology of the TF-gene interaction network into consideration and yield results that are purely based on the local overlap of TF targets and the set of DEGs. In Fig 6 we observe that many TFs inferred by our method, have low enrichment scores.

**Fig 6.**
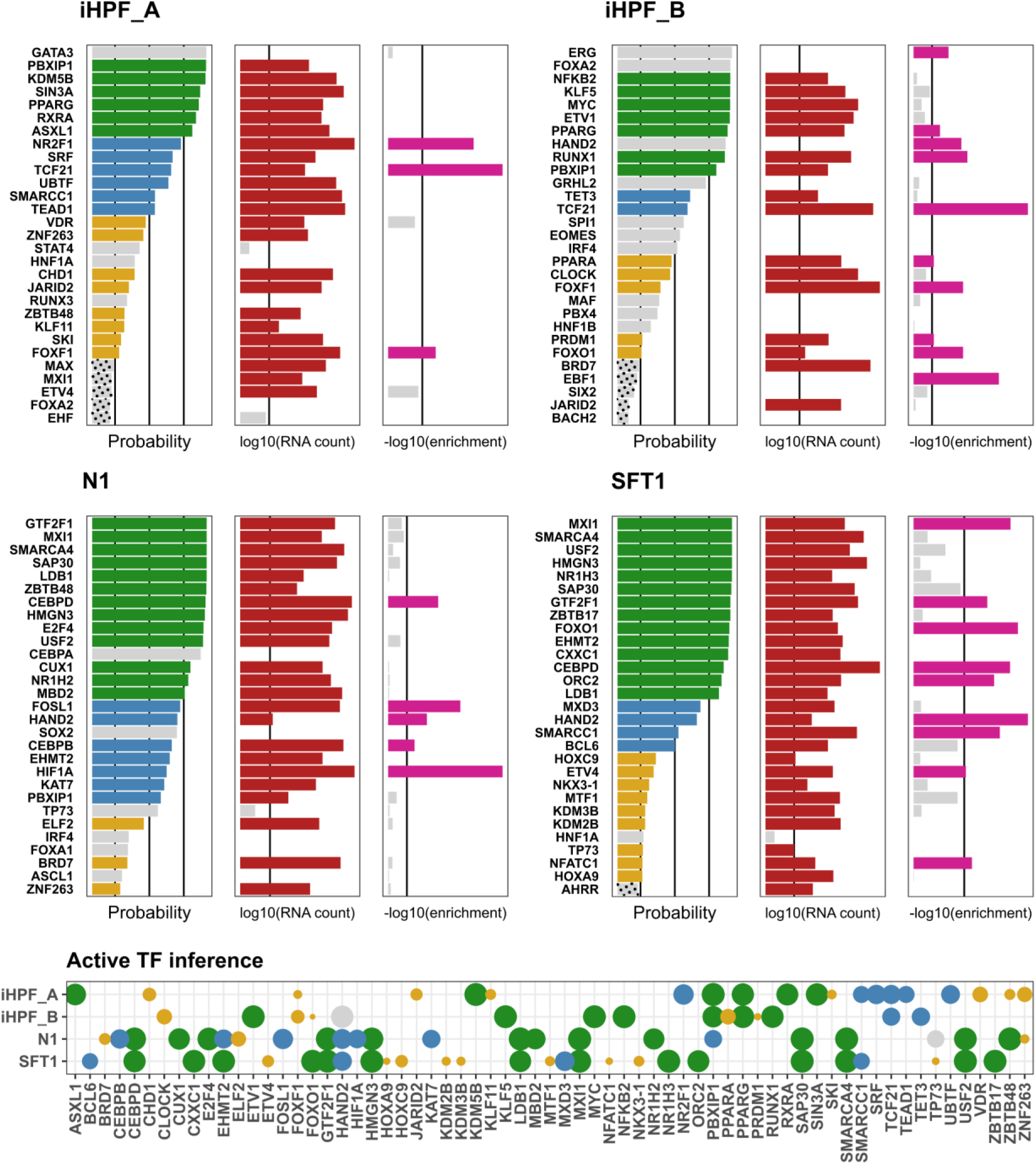
Active TF inference for each cluster of immortalized cells. Gene markers were identified by comparison with pHPF and fed into the inference algorithm. In each data set, the left panel bar plots show the inferred probability of regulator activity by the algorithm. Vertical lines mark 0.2, 0.5, and 0.8 probability thresholds. The middle panel bar plots show the Enrichment Analysis of DE targets (Fisher’s Exact test) for comparison. Significant p-values (<0.05) are highlighted in pink. The right panel bar plots show mean RNA expression across single cells. TFs with expression above 25-percentile are highlighted in red. TFs with inference posterior probability *p* > 0.2, 0.5, 0.8 that also show significant enrichment and RNA expression are highlighted in yellow, blue, and green respectively. TFs with posterior probability *p* > 0.2, but the expression level below the 25 percentile are highlighted in gray. TFs with *p* < 0.2 are shown with a dotted pattern. The bottom panel shows active TFs inferred in each cell line (rows) by the algorithm. N1 and SFT show a similar pattern of TF activity.

Among the TF regulators identified, HAND2 was shared across 2 cell lines. The protein encoded by this gene belongs to the basic helix-loop-helix family of transcription factors and, among many other development-related functions, is required for vascular development and regulation of angiogenesis, possibly through a VEGF signaling pathway (Yamagishi, Olson, & Srivastava, 2000).

The N1 and SFT1 cell lines shared expression of CEBPD, GTF2F1, and MXI1. CEBPD is an intron-less gene that encodes a bZIP transcription factor which can bind as a homodimer to certain DNA regulatory regions. It can also form heterodimers with the related protein CEBP-alpha. The encoded protein is important in the regulation of genes involved in immune and inflammatory responses and may be involved in the regulation of genes associated with activation and/or differentiation of macrophages. It may also be involved in the early stages of adipogenesis (Balamurugan & Sterneck, 2013; Hishida, Nishizuka, Osada, & Imagawa, 2009).

GTF2F1 encodes TFIIF, a general transcription initiation factor that binds to RNA polymerase II and helps to recruit it to the initiation complex in collaboration with TFIIB. It is also a JNK1/3-binding partner and may modulate c-JUN-mediated MAPK signaling in cell proliferation, differentiation, migration, senescence, and apoptosis (Sun, et al., 2015). MXI1 encodes a basic helix-loop-helix protein that inhibits the transcriptional activity of MYC by sequestering MAX, thus preventing the formation of MYC-MAX heterodimers, and by competing with MYC-MAX heterodimers for binding to target sites (Zervos, Gyuris, & Brent, 1993).

The iHPF_A and iHPF_B cell lines shared expression of the TF regulators PPARG and TCF21. PPARG encodes a member of the peroxisome proliferator-activated receptor (PPAR) subfamily of nuclear receptors. PPARs form heterodimers with retinoid X receptors (RXRs) and these heterodimers regulate transcription of various genes that regulate adipocyte differentiation and, pathologically, the development or progression of obesity, diabetes, atherosclerosis and cancer (Rosen, 2005; Cataldi, Costa, Ciccodicola, & Aprile, 2021). TCF21 encodes a transcription factor of the basic helix-loop-helix family. The TCF21 product is mesoderm specific, and expressed in embryonic epicardium, mesenchyme-derived tissues of lung, gut, gonad, and both mesenchymal and glomerular epithelial cells in the kidney. It is involved in the differentiation of mesenchymal cells to fibroblasts (Lighthouse & Small, 2016).

Of note, many of these transcriptional regulators are basic helix-loop-helix TFs, and three TF regulators in particular – CEBPD, TCF21, and HAND2 have been identified as promotors of mesenchymal cell differentiation towards the fibroblast lineage (as opposed to the smooth muscle cell lineage). This suggests that the immortalized fibroblast cell lines express TF regulators that function to maintain the fibroblast phenotype as well as those that may extend this phenotype towards that of immune/inflammatory cells (CEBPD), adipocytes (CEBPD, PPARG) or vascular cells (HAND2). This suggests that fibroblast phenotypic plasticity is perhaps a common rather than exceptional cellular state that may be identified by the expression of particular TF regulators.

## Discussion

In this work, we presented an algorithm for inference of TF activity from differential gene expression profiles and causal graphs. The algorithm incorporates transcriptional logic in the context of Bayesian networks, allowing for probabilistic deviation from deterministic logic rules. The probabilistic framework provides the flexibility for ‘plug-and-play’ integration of various logic models. In this study, we focused on one such model (OR-NOR logic). As a future direction we plan to extend the packages so users can choose the logic prior to running the inference.

The queries are run on causal graphs of TF-gene interactions. We provide several options for such graphs assembled from small-scale curated databases (Kolchanov, et al., 2002; Han, et al., 2015), large-scale public databases (Kanehisa & Goto, 2000; Jensen, et al., 2009; Cerami, et al., 2011), as well as *de novo* reconstructed graphs from high-throughput experiments (Farahmand, O’Connor, Macoska, & Zarringhalam, 2019). We note that the quality and the coverage of the causal graph has a major impact on the ability of regulator activity inference models. Most curated publicly available network of transcriptional regulation with annotation on mode of regulation are small and very limited in their coverage while other higher coverage networks may consist of noisy inferred interactions. Unlike standard enrichment analysis methods, our framework has been designed to account for noise (applicability of interactions and noise in direction of regulation).

Bayesian Networks are Directed Acyclic Graphs (DAGs) and as such, feedback loops cannot be directly modeled in this context, which is a limitation of this approach. Another limitation of our approach is that it is designed to detect TF activation, but not TF deactivation. Moreover, since we only consider the OR-NOR transcriptional regulatory logic, results produced by this approach may miss TFs with alternative regulatory relationships. Since the approach is Bayesian and takes the entire topology of the network into account, by design it outputs a minimal number of TFs whose activation can explain the gene expression data. This is an advantage of our algorithm over enrichment analysis methods that typically contain a large proportion of false positive. The disadvantage may be that sometimes not all true positives get high posteriors probabilities, especially if there are many active regulators present.

For the inference process, we utilized Gibbs Sampling, an MCMC algorithm that is widely used in Bayesian networks. A drawback from MCMC models in Bayesian networks is the convergence time. We implemented the core of the inference in C++ to reduce the wait time. Other strategies can be taken to speed up processing time. For instance, an enrichment-based test can be run a priori to exclude TFs with insufficient differentially expressed targets. This will result in a significant speed up in convergence time, albeit some border line cases may be lost.

Our tool is an exploratory discovery tool that provides a narrow list of potentially relevant TFs, summarizing the observed differential gene expression data. This is similar to standard GO term and pathway enrichment analysis that are also typically applied to summarize differential gene expression data. The focus of our tool is transcriptional regulation and our algorithm can be used as a complementary tool in conjunction with enrichment analysis methods.

To increase the utility of our algorithm, we provided a user-friendly R and python packages as well as a web-based platform with integrated interactive visualization. The pre-processing steps for speeding up the algorithm are implemented as default in the webserver. As databases of causal transcriptional regulatory interactions become more available, we will integrate them in the web-platform and accordingly optimize the inference algorithm for each network.

## Materials and Methods

### Noisy Logic-based gene regulation graphical model

As a starting point, we construct a causal graph from the TF-gene interaction network as follows. The causal graph is a triplet (*G*, *E*, *S*), where *G* represent the nodes, *E* represent the edges (pairs of nodes) and *S* represent signs associated with each edge (+, −). Figure 7 shows a graphical representation of the proposed model. The nodes in the graph consist of the following layers:

- Transcript nodes *Y* = {*Y*_1_, …, *Y*_*m*_}: These are the terminal nodes in the network and represent the transcripts. The domain of these nodes is *D*(*Y*) = {(−), (0), (+)}, representing *downregulated, not regulated, and upregulated* respectively. The value of these nodes will be populated from the input gene expression data.
- True states nodes *H* = {*H*_1_, ⋯, *H*_*m*_}: These nodes represent the true unobserved state of the transcript nodes. This is done to account for noise in input data. These nodes have domain *D*(*H*) = *D*(*Y*) and are central in the implementation of noisy logic gates.
- Regulator state nodes *X* = {*X*_1_, ⋯, *X*_*n*_}: These nodes represent the activation state of TFs in the network. Here we use *D*(*X*) = {(0), (+)}, for no activation or activation respectively.
- TF activity noise nodes θ = {θ_1_, ⋯, θ_*n*_}. For each *X*_*i*_ in the network, we assign a node θ_*i*_, representing a continuous random variable with domain *D*(θ_*i*_) = [0,1]. These nodes represent the probability of activation for the corresponding node *X*_*i*_ and are modeled by a beta distribution.
- Mode of regulation *S* = {*S*_11_, …, *S*_1*m*_, …, *S*_*ij*_, …, *S*_*nm*_}. These nodes represent the mode of action (activation vs. repression) between parent TF node *X*_*i*_ and target transcript node *H*_*j*_. These nodes have a domain *D*(*S*) = {(1), (*NA*), (*A*)}, representing inhibition (I), non-applicable (NA) and activation (A) respectively. We use one-hot encoding for this variable, i.e., *S* is a vector of size 3 with components 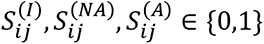.

**Fig 7.**
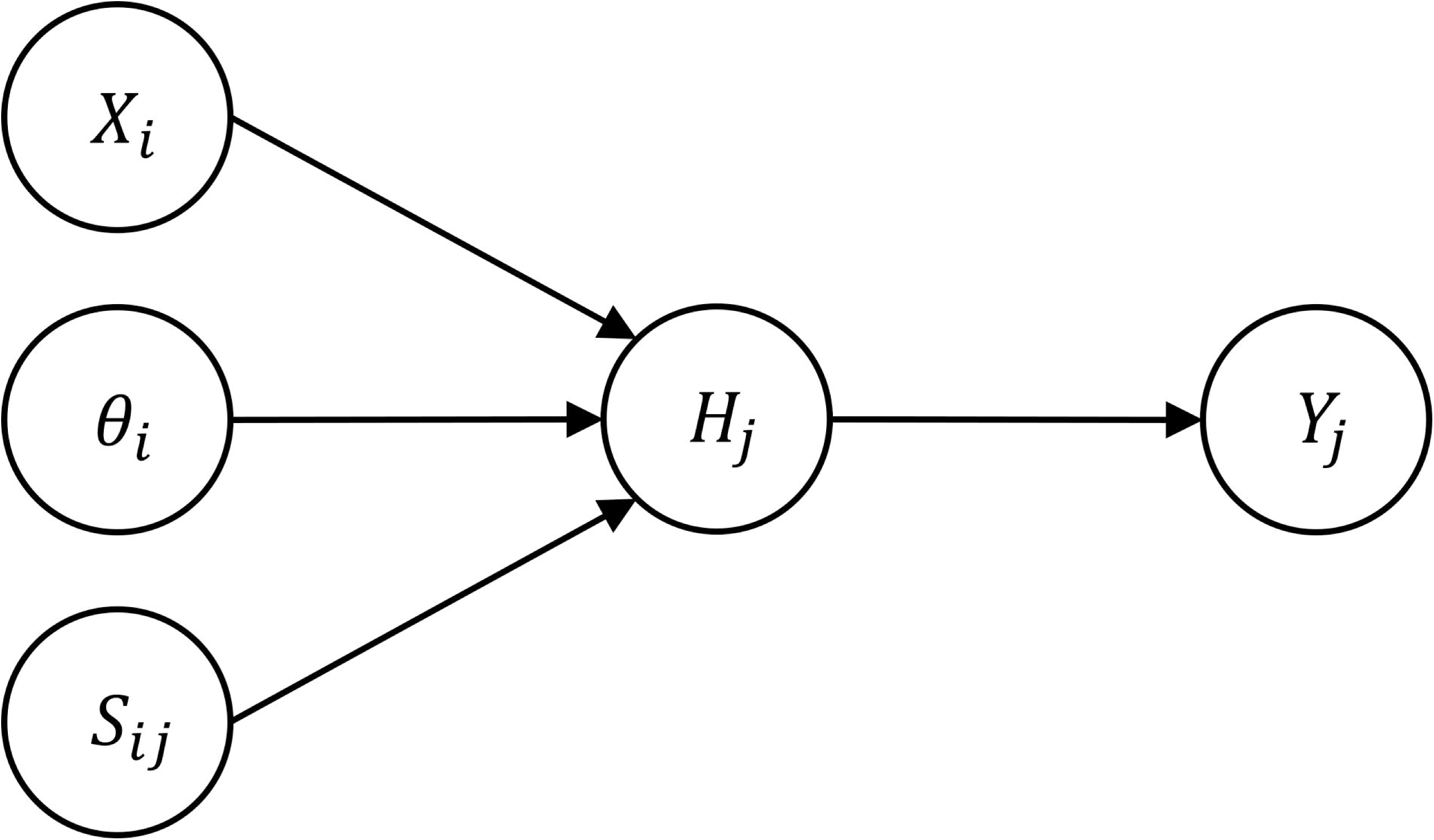
The proposed graphical model for each interaction *i* → *j*.

### Transcriptional Logic

In our model, we incorporate logic gates as in (Bulashevska & Eils, 2005) to explicitly account for combinatorial effects using Boolean logic while accounting for uncertainty. In this work, we consider a combination of noisy OR and NOR gates. We consider two models as follows.

### OR model

This model is used to describe the likelihood of downregulation of a gene by a set of TFs. In this model, presence of one active inhibitor is sufficient to downregulate the gene. The probability mass function is modeled as a Bernoulli trial with probability of success

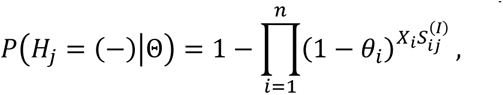

where Θ = {*X*, θ, *S*} all model parameters involving *H* nodes. Although this model seems like a sensible choice, it assumes that all the genes targeted by a TF should strictly follow its influence. However, target regulation depends on many other factors, and we should expect only a fraction of targets to be effectively regulated by a given TF. To make our model more flexible, we incorporate a hyper-parameter ξ, that allows the likelihood model to tolerate more zero-genes in the evidence data. The OR model above is now

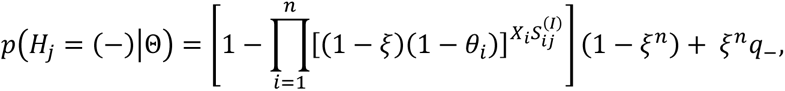

where *q*_−_ represents a prior probability of finding a downregulated gene in the evidence data. For convenience, let us define ζ = (1 − ξ), and let ζ^*deg*^ and ζ^*nnonn*−*deg*^ denote two different values for the ζ parameter that are set depending on whether the current gene has been observed as a modulated or not, respectively. The value of the parameter ζ is set close but greater than zero, for non-differentially expressed genes, e.g. 0 < ζ^*nnonn*−*deg*^ ≤ 0.1, while for differentially expressed genes it may be set close or equal to one. Additionally, an extra (1 − ξ) term is now multiplying (1 − θ_*i*_), effectively increasing the sensitivity of these TF-gene interactions. This has proven beneficial to improve specificity of the inference results. We set ζ^*deg*^ = 0.99 and ζ^*nnonn*−*deg*^ is set to be proportional to *N*_*edges*→*deg*_, i.e., the number of edges in the network that point to genes that are observed as differentially expressed. More specifically, we have used the following relation:

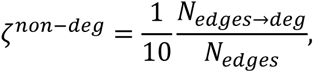

where *N*_*edges*_ is the total number of edges in the network.

### OR-NOR model for gene activation

This model offers a relatively simple way for describing combinatorial effects of both up- and down-regulation within the same interaction network. The rationale is that for a target gene to be activated, at least one of its upstream activators must be activated (OR gate), while at the same time **none** of its inhibitors is active (NOR gate). In this case the target gene up-regulation event is modeled as a Bernoulli trial with probability of success

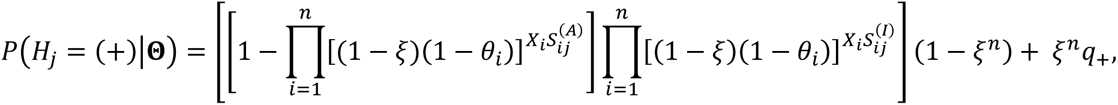

where *q*_+_ is the prior probability of finding an upregulated gene in the evidence data. Target gene activation state is regarded as a multinomial trial with three possible outcomes: upregulation, downregulation, or not changed. This is congruent with discretized differential expression data and allows building a complete model likelihood. Correspondingly, the complementary outcome likelihood is then represented by a NOR-NOR model.

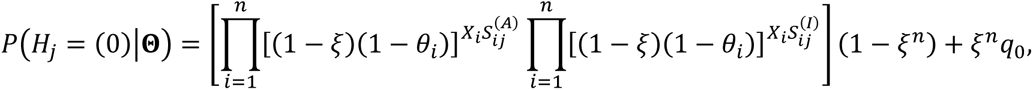

Here *q*_0_ is the prior probability of finding a non-differentially expressed gene in the observed data.

### The model likelihood

The posterior probability of model parameters given the observed data is given by:

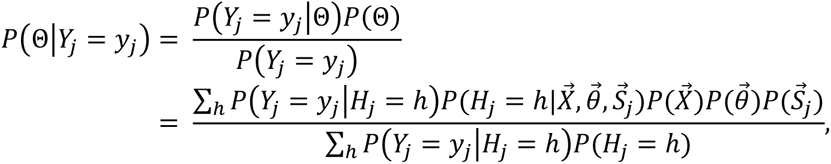

where *P*(*Y*|*H*) is the conditional probability of the observed expression value *Y* given the true state. This conditional probability models the true positive and false positive rate in input gene expression data. **Table 3** shows the values used in our implementation. These values are estimations based on typical experimental errors. This component of the model may encompass several sources of uncertainty, such as dropped reads during RNA sequencing or type I errors in the statistical analysis made for computation of differential gene expression. This may be treated as a prior probability, representing our belief that we would observe these rates.

**Table 3.** Conditional probability of False positive and False negative in the true state given the observed state.

| $P(Y = \cdot \vert H = \cdot)$ | $H = (-)$ | $H = (0)$ | $H = (+)$ |
| --- | --- | --- | --- |
| $Y = (-)$ | 0.945 | 0.050 | 0.005 |
| $Y = (0)$ | 0.050 | 0.900 | 0.050 |
| $Y = (+)$ | 0.005 | 0.050 | 0.945 |

### Fitting the model

The next task is to find the set of model parameters that maximize the model likelihood. Given the large scale of the parameter space, this problem is intractable analytically. A widely used approach for inferring posterior probability in large scale Bayesian networks is Markov Chain Monte Carlo (MCMC) sampling. In particular, Gibbs sampling is a suitable MCMC method to approximate the posterior distribution of the model parameters given the observed data. In Gibbs sampling, we sequentially sample from each random variable, conditioned on the current state of its Markov blanket. To assess the convergence to the posterior distribution of the model parameters, we run at least three independent sampling chains and periodically compute the Gelman-Rubin statistic for each random variable (Gelman & Rubin, 1992). We stop the process after this diagnostic statistic is less than 1.1 for every random variable in the model. The core of the sampling algorithm has been implemented using C++, and user-friendly R and python packages developed.

### Algorithm

In this section we present **Algorithm 1**, which was used throughout this work, along with a detailed description of each of the steps.

#### Algorithm 1.

Sample model’s posterior distribution

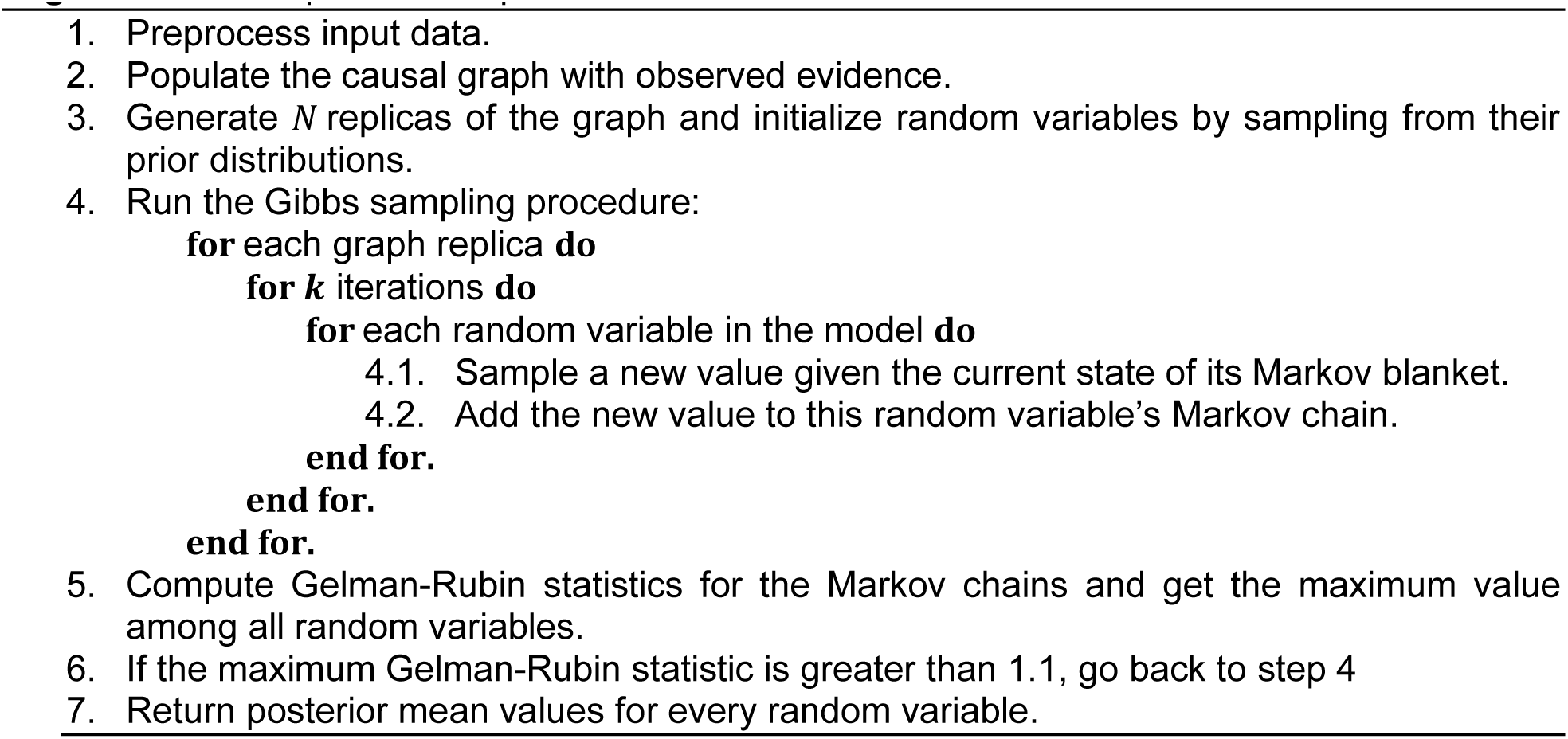

Step 1: Data preprocessing. The input differential gene expression data should contain the computed p-values and fold-change scores for all genes. In this step, we select thresholds for p-values and fold change. The aim is to limit the input data to include only the most significant DEGs, totaling to <= 800 as a rule of thumb.
Step 2: Populate the causal graph. Here we assign the observed values (evidence) for the differential expression of each gene to the corresponding *Y* nodes in the graph. These values will remain fixed during the sampling process.
Step 3: We create *N* independent copies of the graph, allowing us to store separate states of *N* Markov chains to sample. The use of multiple Markov chains provides a way to compare independent results and enables early stopping. Ideally, all *N* chains should converge to the same posterior distribution for all random variables in the model.
Step 4: Gibbs sampling. For each random variable we retrieve the current state of its Markov blanket, which is given by the current values of the children nodes, the parent nodes, and the children of the parent nodes. Then, given this Markov blanket, we compute the conditioned probability of the random variable, from which a new sample is drawn. The sampled value is stored and we move the next random variable and repeat the process. After completing the sampling process for all variables, we start over from the first random variable. This process is repeated for each of the *N* graph replicas created in Step 3. To assess convergence of the posterior distribution, we pause the sampling process after completing *k* rounds to compute the Gelman-Rubin statistics described in the next Step of the algorithm. The choice of k, the period in which convergence is checked, is arbitrary and has been set to *k* = 20 in our implementation.
Step 5: Maximum Gelman-Rubin statistic computation. During the Markov chains sampling process, we need to determine whether the sampled distributions for the random variables in the model have converged to their respective posterior distributions. For this purpose, we compare the *N* Markov chain distributions for every single variable in the model by means of the Gelman-Rubin statistic *R*, which is a dimensionless score that combines the between chain and within chain variances of the random variables. The statistic approaches *R* = 1 as the sampled distributions converge to a steady state. This score is computed for all random variables and we keep the maximum value obtained.
Step 6: Convergence assessment. Convergence of the sampling process is called when max (*R*) < 1.1 for every random variable. If this condition has not been satisfied, we continue the sampling process in Step 4.
Step 7: At this point the sampling process is complete, and the mean value for each random variable is returned.

### Simulations

To assess the impact of noise on model performance, we simulated interaction networks and input differential gene expression data. First, a random interaction network was generated by selecting a total number of *N*_*X*_ TFs and a total number of *N*_*Y*_ genes in the network. For each TF, its number of target genes was sampled according to a negative binomial distribution. Target genes were assigned randomly using a uniform distribution among all genes in the network. Finally for each interaction edge a sign of regulation was assigned with 0.65 probability of being activation (+) and 0.35 probability for repression (-). Subsequently differential gene expression data was simulated as follows. First, we select a random set of TFs that are assigned as *active.* This set of TFs constitute the ground truth. For each active TF, we select 10% of its target genes and assign differential expression according to the sign of regulation as predicted by the graph: +1 if target is upregulated by TF, -1 if downregulated. For genes targeted by multiple active TFs, we perform the algebraic sum of all incoming interactions and take the net sign.

### TF-gene interaction network

The TF-gene interaction network was obtained from (Farahmand, O’Connor, Macoska, & Zarringhalam, 2019) in which interaction network were constructed from direct experimental evidence, integrating data from ChIP-Atlas (Wang, et al., 2018) and The Genotype-Tissue Expression (GTEx) databases (Oki, et al., 2018; Consortium, et al., 2017). Integration was achieved through a regularized Gaussian Graphical model that softly integrated TF-gene interactions derived from ChIP-Seq data into gene expression derived from tissues, resulting in 15 tissue specific TF-gene interaction networks as well as a “merged network” obtained by overlapping tissue-specific networks. In the present work we use the merged network containing interactions that are consistent in at least three tissue types, resulting in 338680 TF-gene interactions from 750 TF molecules.

### Differential gene expression data

DNA microarray-based gene expression profiles for over-expression studies in human primary mammary epithelial cell cultures (Bild, et al., 2006), were obtained from the GEO repository (GSE3151) by using the GEO2R tool for sample selection (Barrett, et al., 2012). Differential expression was computed using the limma R package (Smyth, 2005). We limited the number of differentially expressed genes by applying cutoff thresholds for the adjusted p-values (*p* ≤ 0.01) and log2-foldchange (*fc* ≥ 1) for E2F3 and MYC datasets. For the RAS dataset these cutoff values produced 2226 differentially expressed genes. To further limit the number of DEGs in this experiment, we increased the log2-foldchange cutoff threshold to 2.

Additionally, we used data from a study on fibroblast-to-myofibroblast phenotypic conversion in response to pro-fibrotic signaling molecules TGFβ and CXCL12 (Patalano, et al., 2018; Gharaee-Kermani, et al., 2012; Rodríguez-Nieves, Patalano, Almanza, Gharaee-Kermani, & Macoska, 2016). Differential expression for TGFβ and CXCL12 treated fibroblasts were generated using the R package edgeR (Robinson, McCarthy, & Smyth, 2009).

### Single cell RNA sequencing

Following Trypsin digestion, cells were collected in a 50mL conical tube and washed and resuspended in PBS to a final concentration of ∼700 cells/uL. After gentle resuspension, 2.3ul (∼1,610 cells) per sample was combined with nuclease free water and master mix per 10X recommendations for a targeted recovery of ∼1000 cells per sample.

The samples were loaded onto a 10X Chromium Chip A (PN230027, deprecated) (N1 and SFT1) or Chip G (PN 2000177) (pHPF and iHPF) and run through the 10X Chromium controller instrument and manufacturer protocols for RNA recovery and library preparation. Briefly, cells were partitioned into individual lipid droplets, lysed to release RNA and tagged with UMIs (unique molecular identifiers) for cell of origin. mRNA was isolated with dT oligo beads then reverse transcribed to DNA, ligated with Illumina-compatible sequencing adapters with multiplex capable barcodes (PN-120262 for N1/SFT1 (deprecated), PN-1000213 for pHPF/iHPF), for sample of origin, and PCR amplified for 13 cycles. N1 and SFT1 samples were prepared with Chromium v2 Library and gel bead kit (PN-120267, deprecated) while pHPF and iHPF were prepared with v3 (PN 1000128).

Following library preparation, samples were assessed by Agilent 2100 Bioanalyzer using High Sensitivity chips and reagents (PN 5067-4626) to confirm a normal size distribution to minimize bias, quantified by qPCR with Illumina adapter compatible primers and Sybr Green (Kapa ROX Low Universal Library Quant kit PN KK4873) and molarity calculated by size-correcting to the bioanalyzer average size. N1 and SFT1 samples were pooled together while pHPF and iHPF were pooled together separately.

The samples were sequenced on a Hiseq 2500 in Rapid Run mode in the CPCT Genomics Core, using paired-end on board clustering (PN PE-402-4002) and sequencing by synthesis (SBS) reagents (PN FC-402-4021). Twenty-eight bases were sequenced in read 1 to capture the UMIs, 8 bases for the single indexes, and 91 bases in Read 2 to capture transcripts, yielding ∼100 million total sequencing reads per sample.

### RNA-Seq alignment

Reads were aligned to human genome version GRCh38 using the 10X cellranger v4.0.0 pipeline (cellranger mkref, and cellranger count) using default parameters. Over 93% of reads were mapped to the genome for all 4 datasets. Mapping rates were 93.6%, 93.3%, 96.3% and 96.3% for N1, SFT1, iHPF and pHPF, respectively. To improve the quality of the data, the resulting count matrices were reanalyzed to force the number of cells accepted, to those with highest UMI counts.

### Single Cell Data Analysis

Count data from single cell alignment were processed using the Seurat R package (Stuart, et al., 2019; Hafemeister & Satija, 2019). Seurat objects were created from the filtered matrices resulting from the alignment step. Low quality cells were filtered out, by removing those with large mitochondria contamination and cells with either too few or too many unique genes or total RNA count. Cell cycle scores were assigned to each cell with the method CellCycleScoring using default parameters and cell cycle genes provided by Seurat package. Datasets N1, SFT1, pHPF and iHPF were combined by using Seurat’s merge function and normalized with SCTransform using 3000 variable features and no centering. Batch effect was removed by considering the number of genes, RNA counts and mitochondrial RNA contamination as unwanted sources of variation. Dimensionality reduction was performed through principal component analysis (PCA) and Uniform Manifold Approximation & Projection (UMAP) as implemented in Seurat package, with functions RunPCA (30 PCs) and RunUMAP respectively. Cells were then filtered to work with G1 cells only. Differential expression was computed with respect to pHPF, by using the FindMarkers method. All Seurat methods were used with default parameters, unless otherwise stated.

Prostate tissue single cell data was retrieved from (Henry, et al., 2018). This corresponds to FACS sorted cells, containing fibroblasts, smooth muscle, endothelial and epithelial cells. Here we ignore FACS sorting labels as we run our own classification process. This dataset was preprocessed using Seurat and its SCTransform pipeline, same as with cell lines data. Cells were clustered by using methods FindNeighbors and FindClusters. Each cluster was classified by looking into cell type markers taken from (Henry, et al., 2018), to assign labels: fibroblast, smooth muscle, endothelia, basal epithelia, luminal epithelia, and other epithelia 1 and 2. Cells in G1 cell cycle phase were retained for downstream analysis. Table 4 shows the markers used for the tissue cells classification.

**Table 4.** Gene markers used for classification of single cells in the prostate tissue single cell dataset by (Henry, et al., 2018). These are the same markers used in that study.

| Tissue type | Gene markers |
| --- | --- |
| Basal Epithelia | KRT14, DST, KRT15, KRT5, RGCC |
| Luminal Epithelia | MSMB, KLK3, ACPP, PLA2G2A, KLK2 |
| Other Epithelia 1 | SCGB3A1, LCN2, PIGR, WFDC2, FCGBP |
| Other Epithelia 2 | KRT13, APOBEC3A, CSTB, LYPD3, SERPINB1 |
| Fibroblast | APOD, FBLN1, PTGDS, CFD, DCN |
| Smooth muscle | TPM2, ACTA2, RGS5, MT1A, MYH11 |

Cell lines data and tissue data were combined by using Seurat’s integration pipeline with SCTransform, using 3000 features, k.anchor = 6, and reduction = ‘rpca’.

## Acknowledgements

This work was supported by grants AI150090 (KZ), R01AI167570 (KZ), DK104310 (JAM), and CA156734 (JAM, KZ) from the National Institutes of Health. AA was supported in part by College of Science and Mathematics Dean’s Doctoral Research Fellowship through fellowship support from Oracle, project ID R20000000025727. The funders had no role in study design, data collection and interpretation, or the decision to submit the work for publication.

## Data and software availability

“scRNAseq of Primary and Immortalized Human Prostate Fibroblast Cell Lines” BioProject Accession number: PRJNA881605

Study Accession number: SRP397809

SRA Accession numbers: SRX17617080, SRX17617081, SRX17617082, SRX17617083 Use NCBI’s SRA toolkit to download the 4 datasets above. For further instructions, see: https://www.ncbi.nlm.nih.gov/sra/docs/sradownload

We make our inference algorithm available to use through the following web application: https://umbibio.math.umb.edu/nlbayes

Open-source R and Python packages are available at Github: https://github.com/umbibio/nlbayes-r (doi:10.5281/zenodo.7105306) https://github.com/umbibio/nlbayes-python (doi:10.5281/zenodo.7105233)

We have used R version 4.1.3 and Python 3.10 to develop and test the corresponding packages. Detailed instructions and examples are available in each corresponding repository.

## Supporting Information Legend

1. Supplementary Materials
  a. SI File: SI_File.pdf. Contains additional information, supplementary tables and supplementary figures as follows:
    i. SI File Table S1. Cell lines used in this study.
    ii. SI File Table S2. Top 10 expressed genes in each cell line.
    iii. SI File Table S3. Top 10 differentially expressed genes in each cell line using pHPF as background.
    iv. SI File Fig S1. A) Total number of DEGs compared to the background model (pHPF). B) GO term Enrichment analysis of down regulated genes in each cell line (columns).
    v. SI File Table S4. Simulation performance for several regulatory network configurations. AUC scores are computed on the aggregated results for 30 replicas. Table span is 5 pages.
  b. SI Table 1: SI_Table_1.xlsx. Contains list of Differentially expressed genes in each cell line compared to background pHPF.

