## Supplementary material for "A Bayesian Noisy Logic Model for Inference of Transcription Factor Activity from Single Cell and Bulk Transcriptomic Data": SI File

**Table S1.** Cell lines used in this study.

| Cell line name | Description |
|---|---|
| N1 | 1st line developed in Makosca lab grown from a stromal Nodule of benign prostatic hyperplasia. |
| SFT1 | 1st line grown from a prostatic Spontaneous Fibrous Tumor. |
| pHPF | Primary Human Prostate Fibroblast |
| iHPF | Immortalized Human Prostate Fibroblast |

**Table S2.** Top 10 expressed genes in each cell line.

| | iHPF_A | iHPF_B | pHPF | N1 | SFT1 |
|---|---|---|---|---|---|
| 1 | MT2A | MT2A | MT2A | MT2A | MT2A |
| 2 | MALAT1 | FTL | VIM | FTH1 | MALAT1 |
| 3 | LGALS1 | MALAT1 | MALAT1 | FTL | FTL |
| 4 | FTL | FTH1 | ACTB | MALAT1 | FTH1 |
| 5 | ACTB | LGALS1 | LGALS1 | LGALS1 | LGALS1 |
| 6 | VIM | ACTB | FTL | VIM | VIM |
| 7 | ANXA2 | ANXA2 | ANXA2 | SNHG5 | ACTB |
| 8 | FTH1 | VIM | FTH1 | ACTB | COL1A1 |
| 9 | COL1A1 | TPM2 | TUBA1B | H2AFZ | SNHG5 |
| 10 | TPM2 | NEAT1 | HSP90AA1 | ANXA2 | NEAT1 |

**Table S3.** Top 10 differentially expressed genes in each cell line using to pHPF as background.

| | iHPF_A | iHPF_B | N1 | SFT1 |
|---|---|---|---|---|
| 1 | BEX1 | PLAC8 | SAA1 | SAA1 |
| 2 | EREG | LINC02577 | PITX1 | CHI3L1 |
| 3 | MYEOV | TNFRSF11B | GREM1 | PITX1 |
| 4 | WNT5A | PITX1 | STC2 | RPS29 |

| | | | | |
|---|---|---|---|---|
| **5** | MT1E | SLC4A4 | AKR1B1 | ATP5ME |
| **6** | IGFBP5 | ANGPT1 | ATP5ME | NDUFB1 |
| **7** | TGM2 | IGFBP5 | CHI3L1 | GREM1 |
| **8** | GREM1 | F3 | CDKN2A | CDKN2A |
| **9** | IFI6 | ADAMTS1 | CXCL1 | HIST1H4C |
| **10** | KRTAP2-3 | DIO2 | AREG | AREG |

Fig S1. A) Total number of DEGs compared to the background model (pHPF). B) GO term Enrichment analysis of down regulated genes in each cell line (columns).

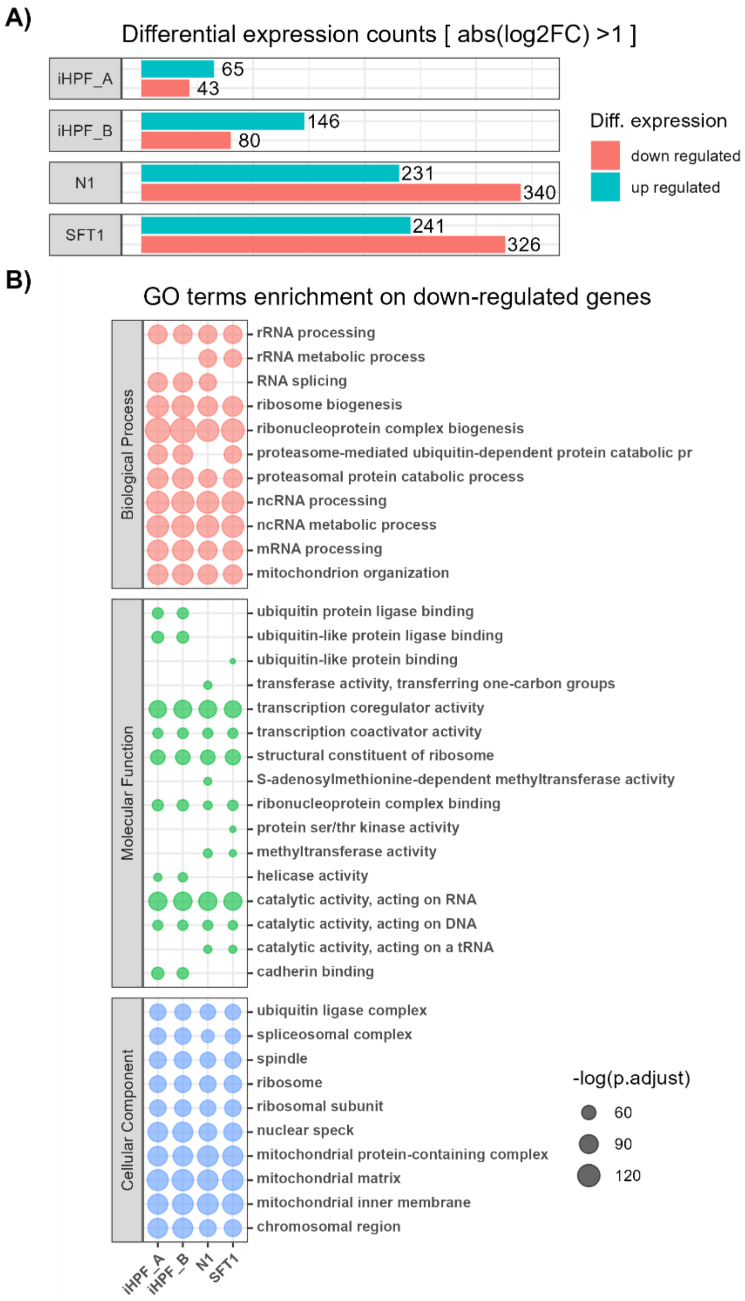

**Table S4.** Simulation performance for several regulatory network configurations. AUC scores are computed on the aggregated results for 30 replicas. Table span is 5 pages.

| N TFs | N Genes | N Edges (approx.) | Up Reg. | Down Reg. | N Act. TFs | N DEG (avg) | No Noise ROC AUC | No Noise PRC AUC | 25% Noise ROC AUC | 25% Noise PRC AUC |
|---|---|---|---|---|---|---|---|---|---|---|
| 250 | 5000 | 16650 | 35% | 65% | 10 | 55.1 | 0.996 | 0.983 | 0.976 | 0.877 |
| 250 | 5000 | 16650 | 35% | 65% | 30 | 172.3 | 0.990 | 0.969 | 0.942 | 0.836 |
| 250 | 5000 | 16650 | 35% | 65% | 50 | 285.9 | 0.983 | 0.959 | 0.919 | 0.813 |
| 250 | 5000 | 16650 | 50% | 50% | 10 | 55.6 | 0.996 | 0.984 | 0.971 | 0.872 |
| 250 | 5000 | 16650 | 50% | 50% | 30 | 172.1 | 0.990 | 0.974 | 0.948 | 0.850 |
| 250 | 5000 | 16650 | 50% | 50% | 50 | 284.3 | 0.984 | 0.962 | 0.925 | 0.828 |
| 250 | 5000 | 16650 | 65% | 35% | 10 | 55.4 | 0.997 | 0.986 | 0.970 | 0.869 |
| 250 | 5000 | 16650 | 65% | 35% | 30 | 173.5 | 0.987 | 0.969 | 0.943 | 0.832 |
| 250 | 5000 | 16650 | 65% | 35% | 50 | 283.6 | 0.983 | 0.959 | 0.917 | 0.817 |
| 250 | 5000 | 33300 | 35% | 65% | 10 | 116.6 | 1.000 | 0.994 | 0.982 | 0.906 |
| 250 | 5000 | 33300 | 35% | 65% | 30 | 359.4 | 0.994 | 0.982 | 0.967 | 0.886 |
| 250 | 5000 | 33300 | 35% | 65% | 50 | 580.4 | 0.989 | 0.975 | 0.931 | 0.838 |
| 250 | 5000 | 33300 | 50% | 50% | 10 | 117.9 | 0.997 | 0.990 | 0.985 | 0.915 |
| 250 | 5000 | 33300 | 50% | 50% | 30 | 357.9 | 0.997 | 0.989 | 0.969 | 0.896 |
| 250 | 5000 | 33300 | 50% | 50% | 50 | 578.9 | 0.993 | 0.981 | 0.940 | 0.856 |
| 250 | 5000 | 33300 | 65% | 35% | 10 | 116.9 | 0.999 | 0.993 | 0.984 | 0.910 |
| 250 | 5000 | 33300 | 65% | 35% | 30 | 360.0 | 0.996 | 0.986 | 0.967 | 0.890 |
| 250 | 5000 | 33300 | 65% | 35% | 50 | 576.0 | 0.990 | 0.976 | 0.937 | 0.850 |
| 250 | 5000 | 49950 | 35% | 65% | 10 | 173.8 | 0.999 | 0.994 | 0.994 | 0.944 |
| 250 | 5000 | 49950 | 35% | 65% | 30 | 524.7 | 0.997 | 0.988 | 0.974 | 0.898 |
| 250 | 5000 | 49950 | 35% | 65% | 50 | 833.8 | 0.985 | 0.967 | 0.934 | 0.836 |
| 250 | 5000 | 49950 | 50% | 50% | 10 | 175.0 | 1.000 | 0.997 | 0.994 | 0.944 |
| 250 | 5000 | 49950 | 50% | 50% | 30 | 524.0 | 0.995 | 0.987 | 0.976 | 0.913 |
| 250 | 5000 | 49950 | 50% | 50% | 50 | 829.9 | 0.987 | 0.972 | 0.938 | 0.845 |
| 250 | 5000 | 49950 | 65% | 35% | 10 | 175.9 | 1.000 | 0.997 | 0.994 | 0.952 |
| 250 | 5000 | 49950 | 65% | 35% | 30 | 522.3 | 0.997 | 0.988 | 0.975 | 0.918 |
| 250 | 5000 | 49950 | 65% | 35% | 50 | 832.5 | 0.989 | 0.974 | 0.939 | 0.850 |
| 250 | 10000 | 33300 | 35% | 65% | 10 | 118.6 | 1.000 | 0.999 | 0.990 | 0.945 |
| 250 | 10000 | 33300 | 35% | 65% | 30 | 369.9 | 0.999 | 0.996 | 0.978 | 0.934 |
| 250 | 10000 | 33300 | 35% | 65% | 50 | 607.6 | 0.997 | 0.993 | 0.961 | 0.913 |
| 250 | 10000 | 33300 | 50% | 50% | 10 | 119.4 | 1.000 | 0.997 | 0.987 | 0.937 |
| 250 | 10000 | 33300 | 50% | 50% | 30 | 367.5 | 0.998 | 0.994 | 0.983 | 0.941 |
| 250 | 10000 | 33300 | 50% | 50% | 50 | 605.3 | 0.997 | 0.993 | 0.971 | 0.930 |
| 250 | 10000 | 33300 | 65% | 35% | 10 | 119.5 | 0.999 | 0.996 | 0.986 | 0.927 |
| 250 | 10000 | 33300 | 65% | 35% | 30 | 366.1 | 0.998 | 0.994 | 0.973 | 0.922 |
| 250 | 10000 | 33300 | 65% | 35% | 50 | 604.1 | 0.996 | 0.991 | 0.964 | 0.922 |
| 250 | 10000 | 66600 | 35% | 65% | 10 | 242.0 | 0.999 | 0.995 | 0.997 | 0.969 |
| 250 | 10000 | 66600 | 35% | 65% | 30 | 730.9 | 1.000 | 0.998 | 0.990 | 0.961 |
| 250 | 10000 | 66600 | 35% | 65% | 50 | 1184.3 | 0.997 | 0.995 | 0.971 | 0.925 |
| 250 | 10000 | 66600 | 50% | 50% | 10 | 242.7 | 1.000 | 0.997 | 0.996 | 0.966 |
| 250 | 10000 | 66600 | 50% | 50% | 30 | 730.6 | 0.999 | 0.998 | 0.990 | 0.962 |
| 250 | 10000 | 66600 | 50% | 50% | 50 | 1178.1 | 0.998 | 0.996 | 0.979 | 0.945 |
| 250 | 10000 | 66600 | 65% | 35% | 10 | 240.9 | 1.000 | 0.998 | 0.994 | 0.960 |
| 250 | 10000 | 66600 | 65% | 35% | 30 | 731.1 | 1.000 | 0.998 | 0.991 | 0.962 |
| 250 | 10000 | 66600 | 65% | 35% | 50 | 1180.8 | 0.998 | 0.994 | 0.979 | 0.947 |
| 250 | 10000 | 99900 | 35% | 65% | 10 | 362.7 | 1.000 | 1.000 | 0.999 | 0.990 |
| 250 | 10000 | 99900 | 35% | 65% | 30 | 1070.6 | 0.999 | 0.997 | 0.993 | 0.969 |
| 250 | 10000 | 99900 | 35% | 65% | 50 | 1692.9 | 0.994 | 0.991 | 0.979 | 0.946 |

| | | | | | | | | | | |
|---|---|---|---|---|---|---|---|---|---|---|
| 250 | 10000 | 99900 | 50% | 50% | 10 | 362.2 | 1.000 | 1.000 | 0.999 | 0.991 |
| 250 | 10000 | 99900 | 50% | 50% | 30 | 1068.2 | 0.999 | 0.998 | 0.993 | 0.978 |
| 250 | 10000 | 99900 | 50% | 50% | 50 | 1693.0 | 0.995 | 0.992 | 0.982 | 0.951 |
| 250 | 10000 | 99900 | 65% | 35% | 10 | 362.8 | 1.000 | 1.000 | 0.999 | 0.992 |
| 250 | 10000 | 99900 | 65% | 35% | 30 | 1072.7 | 0.999 | 0.998 | 0.995 | 0.977 |
| 250 | 10000 | 99900 | 65% | 35% | 50 | 1699.7 | 0.995 | 0.992 | 0.984 | 0.958 |
| 250 | 15000 | 49950 | 35% | 65% | 10 | 178.7 | 1.000 | 1.000 | 0.993 | 0.950 |
| 250 | 15000 | 49950 | 35% | 65% | 30 | 550.1 | 1.000 | 0.999 | 0.988 | 0.959 |
| 250 | 15000 | 49950 | 35% | 65% | 50 | 905.6 | 0.999 | 0.997 | 0.979 | 0.951 |
| 250 | 15000 | 49950 | 50% | 50% | 10 | 178.5 | 1.000 | 1.000 | 0.996 | 0.963 |
| 250 | 15000 | 49950 | 50% | 50% | 30 | 549.9 | 1.000 | 0.998 | 0.992 | 0.968 |
| 250 | 15000 | 49950 | 50% | 50% | 50 | 903.7 | 1.000 | 0.999 | 0.981 | 0.952 |
| 250 | 15000 | 49950 | 65% | 35% | 10 | 179.9 | 1.000 | 1.000 | 0.990 | 0.948 |
| 250 | 15000 | 49950 | 65% | 35% | 30 | 551.2 | 1.000 | 0.999 | 0.989 | 0.960 |
| 250 | 15000 | 49950 | 65% | 35% | 50 | 906.5 | 1.000 | 0.999 | 0.979 | 0.950 |
| 250 | 15000 | 99900 | 35% | 65% | 10 | 364.8 | 1.000 | 1.000 | 0.999 | 0.988 |
| 250 | 15000 | 99900 | 35% | 65% | 30 | 1106.8 | 1.000 | 0.999 | 0.994 | 0.976 |
| 250 | 15000 | 99900 | 35% | 65% | 50 | 1783.7 | 0.999 | 0.998 | 0.989 | 0.970 |
| 250 | 15000 | 99900 | 50% | 50% | 10 | 365.5 | 1.000 | 1.000 | 0.999 | 0.988 |
| 250 | 15000 | 99900 | 50% | 50% | 30 | 1103.3 | 1.000 | 1.000 | 0.997 | 0.987 |
| 250 | 15000 | 99900 | 50% | 50% | 50 | 1778.1 | 0.999 | 0.998 | 0.990 | 0.972 |
| 250 | 15000 | 99900 | 65% | 35% | 10 | 365.9 | 1.000 | 1.000 | 0.998 | 0.980 |
| 250 | 15000 | 99900 | 65% | 35% | 30 | 1108.2 | 1.000 | 0.999 | 0.996 | 0.983 |
| 250 | 15000 | 99900 | 65% | 35% | 50 | 1781.6 | 0.999 | 0.999 | 0.988 | 0.969 |
| 250 | 15000 | 149850 | 35% | 65% | 10 | 549.2 | 1.000 | 1.000 | 1.000 | 0.995 |
| 250 | 15000 | 149850 | 35% | 65% | 30 | 1626.5 | 1.000 | 1.000 | 0.997 | 0.991 |
| 250 | 15000 | 149850 | 35% | 65% | 50 | 2576.8 | 0.998 | 0.997 | 0.989 | 0.973 |
| 250 | 15000 | 149850 | 50% | 50% | 10 | 550.0 | 1.000 | 0.999 | 1.000 | 0.996 |
| 250 | 15000 | 149850 | 50% | 50% | 30 | 1620.0 | 1.000 | 1.000 | 0.998 | 0.990 |
| 250 | 15000 | 149850 | 50% | 50% | 50 | 2559.2 | 0.997 | 0.997 | 0.991 | 0.977 |
| 250 | 15000 | 149850 | 65% | 35% | 10 | 547.1 | 1.000 | 1.000 | 1.000 | 0.997 |
| 250 | 15000 | 149850 | 65% | 35% | 30 | 1625.2 | 1.000 | 1.000 | 0.997 | 0.989 |
| 250 | 15000 | 149850 | 65% | 35% | 50 | 2578.7 | 0.998 | 0.998 | 0.991 | 0.977 |
| 500 | 5000 | 16650 | 35% | 65% | 10 | 27.4 | 0.994 | 0.949 | 0.963 | 0.800 |
| 500 | 5000 | 16650 | 35% | 65% | 30 | 80.7 | 0.986 | 0.930 | 0.930 | 0.740 |
| 500 | 5000 | 16650 | 35% | 65% | 50 | 131.9 | 0.978 | 0.909 | 0.926 | 0.719 |
| 500 | 5000 | 16650 | 50% | 50% | 10 | 27.5 | 0.993 | 0.945 | 0.952 | 0.785 |
| 500 | 5000 | 16650 | 50% | 50% | 30 | 80.7 | 0.985 | 0.930 | 0.934 | 0.747 |
| 500 | 5000 | 16650 | 50% | 50% | 50 | 132.7 | 0.980 | 0.918 | 0.929 | 0.746 |
| 500 | 5000 | 16650 | 65% | 35% | 10 | 27.4 | 0.996 | 0.949 | 0.965 | 0.764 |
| 500 | 5000 | 16650 | 65% | 35% | 30 | 81.7 | 0.987 | 0.925 | 0.937 | 0.743 |
| 500 | 5000 | 16650 | 65% | 35% | 50 | 133.3 | 0.980 | 0.910 | 0.921 | 0.721 |
| 500 | 5000 | 33300 | 35% | 65% | 10 | 60.5 | 0.999 | 0.982 | 0.983 | 0.851 |
| 500 | 5000 | 33300 | 35% | 65% | 30 | 178.7 | 0.993 | 0.964 | 0.951 | 0.796 |
| 500 | 5000 | 33300 | 35% | 65% | 50 | 289.8 | 0.984 | 0.941 | 0.930 | 0.752 |
| 500 | 5000 | 33300 | 50% | 50% | 10 | 60.0 | 0.997 | 0.983 | 0.969 | 0.832 |
| 500 | 5000 | 33300 | 50% | 50% | 30 | 179.2 | 0.994 | 0.967 | 0.960 | 0.813 |
| 500 | 5000 | 33300 | 50% | 50% | 50 | 291.1 | 0.984 | 0.943 | 0.930 | 0.758 |
| 500 | 5000 | 33300 | 65% | 35% | 10 | 61.1 | 0.991 | 0.971 | 0.966 | 0.817 |
| 500 | 5000 | 33300 | 65% | 35% | 30 | 180.3 | 0.991 | 0.963 | 0.952 | 0.796 |
| 500 | 5000 | 33300 | 65% | 35% | 50 | 288.5 | 0.983 | 0.939 | 0.933 | 0.776 |
| 500 | 5000 | 49950 | 35% | 65% | 10 | 95.0 | 0.999 | 0.992 | 0.977 | 0.872 |
| 500 | 5000 | 49950 | 35% | 65% | 30 | 276.4 | 0.994 | 0.977 | 0.970 | 0.838 |
| 500 | 5000 | 49950 | 35% | 65% | 50 | 441.0 | 0.989 | 0.956 | 0.942 | 0.788 |

| | | | | | | | | | | |
|---|---|---|---|---|---|---|---|---|---|---|
| 500 | 5000 | 49950 | 50% | 50% | 10 | 94.1 | 1.000 | 0.994 | 0.990 | 0.907 |
| 500 | 5000 | 49950 | 50% | 50% | 30 | 274.6 | 0.997 | 0.981 | 0.971 | 0.854 |
| 500 | 5000 | 49950 | 50% | 50% | 50 | 437.7 | 0.990 | 0.962 | 0.948 | 0.803 |
| 500 | 5000 | 49950 | 65% | 35% | 10 | 94.5 | 0.999 | 0.994 | 0.983 | 0.891 |
| 500 | 5000 | 49950 | 65% | 35% | 30 | 274.1 | 0.996 | 0.981 | 0.973 | 0.857 |
| 500 | 5000 | 49950 | 65% | 35% | 50 | 439.2 | 0.989 | 0.955 | 0.944 | 0.788 |
| 500 | 10000 | 33300 | 35% | 65% | 10 | 61.0 | 0.996 | 0.984 | 0.972 | 0.861 |
| 500 | 10000 | 33300 | 35% | 65% | 30 | 182.2 | 0.995 | 0.984 | 0.975 | 0.880 |
| 500 | 10000 | 33300 | 35% | 65% | 50 | 293.5 | 0.991 | 0.975 | 0.956 | 0.843 |
| 500 | 10000 | 33300 | 50% | 50% | 10 | 61.2 | 0.998 | 0.985 | 0.979 | 0.898 |
| 500 | 10000 | 33300 | 50% | 50% | 30 | 180.0 | 0.997 | 0.986 | 0.973 | 0.884 |
| 500 | 10000 | 33300 | 50% | 50% | 50 | 294.8 | 0.994 | 0.979 | 0.960 | 0.862 |
| 500 | 10000 | 33300 | 65% | 35% | 10 | 60.4 | 0.996 | 0.983 | 0.975 | 0.871 |
| 500 | 10000 | 33300 | 65% | 35% | 30 | 182.4 | 0.994 | 0.982 | 0.972 | 0.876 |
| 500 | 10000 | 33300 | 65% | 35% | 50 | 293.5 | 0.992 | 0.973 | 0.951 | 0.843 |
| 500 | 10000 | 66600 | 35% | 65% | 10 | 128.8 | 1.000 | 1.000 | 0.992 | 0.936 |
| 500 | 10000 | 66600 | 35% | 65% | 30 | 378.2 | 0.999 | 0.996 | 0.987 | 0.925 |
| 500 | 10000 | 66600 | 35% | 65% | 50 | 611.3 | 0.996 | 0.989 | 0.970 | 0.890 |
| 500 | 10000 | 66600 | 50% | 50% | 10 | 127.8 | 1.000 | 1.000 | 0.994 | 0.939 |
| 500 | 10000 | 66600 | 50% | 50% | 30 | 378.0 | 1.000 | 0.997 | 0.987 | 0.925 |
| 500 | 10000 | 66600 | 50% | 50% | 50 | 609.9 | 0.998 | 0.992 | 0.976 | 0.912 |
| 500 | 10000 | 66600 | 65% | 35% | 10 | 128.2 | 1.000 | 0.997 | 0.993 | 0.947 |
| 500 | 10000 | 66600 | 65% | 35% | 30 | 379.8 | 0.999 | 0.996 | 0.988 | 0.930 |
| 500 | 10000 | 66600 | 65% | 35% | 50 | 611.5 | 0.998 | 0.991 | 0.979 | 0.914 |
| 500 | 10000 | 99900 | 35% | 65% | 10 | 195.0 | 1.000 | 1.000 | 0.996 | 0.960 |
| 500 | 10000 | 99900 | 35% | 65% | 30 | 568.5 | 1.000 | 0.999 | 0.995 | 0.961 |
| 500 | 10000 | 99900 | 35% | 65% | 50 | 908.6 | 0.999 | 0.993 | 0.984 | 0.926 |
| 500 | 10000 | 99900 | 50% | 50% | 10 | 194.0 | 1.000 | 1.000 | 0.998 | 0.970 |
| 500 | 10000 | 99900 | 50% | 50% | 30 | 569.5 | 1.000 | 0.998 | 0.996 | 0.966 |
| 500 | 10000 | 99900 | 50% | 50% | 50 | 906.1 | 0.998 | 0.993 | 0.986 | 0.938 |
| 500 | 10000 | 99900 | 65% | 35% | 10 | 193.9 | 1.000 | 1.000 | 0.998 | 0.970 |
| 500 | 10000 | 99900 | 65% | 35% | 30 | 568.8 | 1.000 | 0.998 | 0.993 | 0.958 |
| 500 | 10000 | 99900 | 65% | 35% | 50 | 903.3 | 0.998 | 0.992 | 0.986 | 0.936 |
| 500 | 15000 | 49950 | 35% | 65% | 10 | 95.0 | 0.999 | 0.996 | 0.986 | 0.932 |
| 500 | 15000 | 49950 | 35% | 65% | 30 | 284.0 | 0.999 | 0.996 | 0.983 | 0.921 |
| 500 | 15000 | 49950 | 35% | 65% | 50 | 460.5 | 0.997 | 0.990 | 0.973 | 0.902 |
| 500 | 15000 | 49950 | 50% | 50% | 10 | 94.6 | 1.000 | 0.996 | 0.985 | 0.923 |
| 500 | 15000 | 49950 | 50% | 50% | 30 | 282.6 | 1.000 | 0.997 | 0.984 | 0.925 |
| 500 | 15000 | 49950 | 50% | 50% | 50 | 458.1 | 0.997 | 0.991 | 0.974 | 0.905 |
| 500 | 15000 | 49950 | 65% | 35% | 10 | 95.8 | 1.000 | 0.997 | 0.991 | 0.934 |
| 500 | 15000 | 49950 | 65% | 35% | 30 | 285.6 | 0.998 | 0.994 | 0.982 | 0.919 |
| 500 | 15000 | 49950 | 65% | 35% | 50 | 458.9 | 0.998 | 0.991 | 0.972 | 0.902 |
| 500 | 15000 | 99900 | 35% | 65% | 10 | 194.4 | 1.000 | 1.000 | 0.997 | 0.962 |
| 500 | 15000 | 99900 | 35% | 65% | 30 | 577.0 | 1.000 | 0.999 | 0.992 | 0.954 |
| 500 | 15000 | 99900 | 35% | 65% | 50 | 928.1 | 1.000 | 0.998 | 0.989 | 0.945 |
| 500 | 15000 | 99900 | 50% | 50% | 10 | 196.8 | 1.000 | 1.000 | 0.995 | 0.965 |
| 500 | 15000 | 99900 | 50% | 50% | 30 | 576.9 | 1.000 | 0.999 | 0.995 | 0.967 |
| 500 | 15000 | 99900 | 50% | 50% | 50 | 924.1 | 0.999 | 0.997 | 0.989 | 0.953 |
| 500 | 15000 | 99900 | 65% | 35% | 10 | 195.2 | 1.000 | 1.000 | 0.996 | 0.962 |
| 500 | 15000 | 99900 | 65% | 35% | 30 | 575.3 | 1.000 | 0.998 | 0.995 | 0.969 |
| 500 | 15000 | 99900 | 65% | 35% | 50 | 924.9 | 0.999 | 0.997 | 0.989 | 0.950 |
| 500 | 15000 | 149850 | 35% | 65% | 10 | 292.8 | 1.000 | 1.000 | 1.000 | 0.992 |
| 500 | 15000 | 149850 | 35% | 65% | 30 | 855.0 | 1.000 | 1.000 | 0.998 | 0.981 |
| 500 | 15000 | 149850 | 35% | 65% | 50 | 1364.1 | 0.998 | 0.996 | 0.993 | 0.966 |

| | | | | | | | | | | |
|---|---|---|---|---|---|---|---|---|---|---|
| 500 | 15000 | 149850 | 50% | 50% | 10 | 292.5 | 1.000 | 1.000 | 1.000 | 0.991 |
| 500 | 15000 | 149850 | 50% | 50% | 30 | 856.0 | 1.000 | 0.999 | 0.998 | 0.986 |
| 500 | 15000 | 149850 | 50% | 50% | 50 | 1358.5 | 1.000 | 0.998 | 0.993 | 0.971 |
| 500 | 15000 | 149850 | 65% | 35% | 10 | 294.4 | 1.000 | 1.000 | 1.000 | 0.990 |
| 500 | 15000 | 149850 | 65% | 35% | 30 | 855.5 | 1.000 | 0.999 | 0.998 | 0.983 |
| 500 | 15000 | 149850 | 65% | 35% | 50 | 1360.7 | 1.000 | 0.998 | 0.993 | 0.968 |
| 1000 | 5000 | 16650 | 35% | 65% | 10 | 11.4 | 0.999 | 0.856 | 0.909 | 0.566 |
| 1000 | 5000 | 16650 | 35% | 65% | 30 | 32.3 | 0.978 | 0.831 | 0.911 | 0.561 |
| 1000 | 5000 | 16650 | 35% | 65% | 50 | 52.4 | 0.972 | 0.788 | 0.887 | 0.562 |
| 1000 | 5000 | 16650 | 50% | 50% | 10 | 11.9 | 0.999 | 0.856 | 0.932 | 0.613 |
| 1000 | 5000 | 16650 | 50% | 50% | 30 | 32.7 | 0.978 | 0.832 | 0.901 | 0.595 |
| 1000 | 5000 | 16650 | 50% | 50% | 50 | 54.2 | 0.970 | 0.800 | 0.900 | 0.595 |
| 1000 | 5000 | 16650 | 65% | 35% | 10 | 11.8 | 0.999 | 0.865 | 0.911 | 0.610 |
| 1000 | 5000 | 16650 | 65% | 35% | 30 | 33.1 | 0.982 | 0.830 | 0.910 | 0.604 |
| 1000 | 5000 | 16650 | 65% | 35% | 50 | 52.7 | 0.974 | 0.794 | 0.885 | 0.557 |
| 1000 | 5000 | 33300 | 35% | 65% | 10 | 28.1 | 0.991 | 0.915 | 0.949 | 0.677 |
| 1000 | 5000 | 33300 | 35% | 65% | 30 | 84.4 | 0.985 | 0.871 | 0.940 | 0.649 |
| 1000 | 5000 | 33300 | 35% | 65% | 50 | 137.3 | 0.969 | 0.831 | 0.910 | 0.611 |
| 1000 | 5000 | 33300 | 50% | 50% | 10 | 28.5 | 0.992 | 0.918 | 0.953 | 0.732 |
| 1000 | 5000 | 33300 | 50% | 50% | 30 | 83.9 | 0.984 | 0.885 | 0.928 | 0.675 |
| 1000 | 5000 | 33300 | 50% | 50% | 50 | 136.4 | 0.973 | 0.841 | 0.915 | 0.619 |
| 1000 | 5000 | 33300 | 65% | 35% | 10 | 28.1 | 0.989 | 0.914 | 0.955 | 0.687 |
| 1000 | 5000 | 33300 | 65% | 35% | 30 | 84.4 | 0.977 | 0.871 | 0.934 | 0.649 |
| 1000 | 5000 | 33300 | 65% | 35% | 50 | 137.7 | 0.969 | 0.838 | 0.914 | 0.619 |
| 1000 | 5000 | 49950 | 35% | 65% | 10 | 45.8 | 0.995 | 0.948 | 0.967 | 0.767 |
| 1000 | 5000 | 49950 | 35% | 65% | 30 | 135.6 | 0.991 | 0.929 | 0.952 | 0.718 |
| 1000 | 5000 | 49950 | 35% | 65% | 50 | 219.4 | 0.983 | 0.898 | 0.929 | 0.660 |
| 1000 | 5000 | 49950 | 50% | 50% | 10 | 46.0 | 0.992 | 0.953 | 0.972 | 0.789 |
| 1000 | 5000 | 49950 | 50% | 50% | 30 | 134.7 | 0.994 | 0.940 | 0.957 | 0.724 |
| 1000 | 5000 | 49950 | 50% | 50% | 50 | 217.5 | 0.986 | 0.910 | 0.938 | 0.675 |
| 1000 | 5000 | 49950 | 65% | 35% | 10 | 45.2 | 0.994 | 0.954 | 0.972 | 0.801 |
| 1000 | 5000 | 49950 | 65% | 35% | 30 | 136.0 | 0.992 | 0.932 | 0.952 | 0.712 |
| 1000 | 5000 | 49950 | 65% | 35% | 50 | 218.8 | 0.983 | 0.905 | 0.937 | 0.678 |
| 1000 | 10000 | 33300 | 35% | 65% | 10 | 28.0 | 0.997 | 0.961 | 0.975 | 0.790 |
| 1000 | 10000 | 33300 | 35% | 65% | 30 | 85.2 | 0.995 | 0.941 | 0.938 | 0.744 |
| 1000 | 10000 | 33300 | 35% | 65% | 50 | 138.0 | 0.986 | 0.924 | 0.940 | 0.746 |
| 1000 | 10000 | 33300 | 50% | 50% | 10 | 28.6 | 0.996 | 0.957 | 0.949 | 0.758 |
| 1000 | 10000 | 33300 | 50% | 50% | 30 | 84.2 | 0.993 | 0.940 | 0.947 | 0.751 |
| 1000 | 10000 | 33300 | 50% | 50% | 50 | 137.8 | 0.984 | 0.925 | 0.938 | 0.757 |
| 1000 | 10000 | 33300 | 65% | 35% | 10 | 28.4 | 0.996 | 0.958 | 0.951 | 0.777 |
| 1000 | 10000 | 33300 | 65% | 35% | 30 | 84.5 | 0.993 | 0.942 | 0.940 | 0.756 |
| 1000 | 10000 | 33300 | 65% | 35% | 50 | 137.4 | 0.985 | 0.925 | 0.940 | 0.745 |
| 1000 | 10000 | 66600 | 35% | 65% | 10 | 63.4 | 0.998 | 0.980 | 0.974 | 0.850 |
| 1000 | 10000 | 66600 | 35% | 65% | 30 | 187.3 | 0.996 | 0.977 | 0.970 | 0.821 |
| 1000 | 10000 | 66600 | 35% | 65% | 50 | 302.3 | 0.994 | 0.972 | 0.959 | 0.799 |
| 1000 | 10000 | 66600 | 50% | 50% | 10 | 64.0 | 0.997 | 0.982 | 0.979 | 0.864 |
| 1000 | 10000 | 66600 | 50% | 50% | 30 | 187.5 | 0.997 | 0.980 | 0.975 | 0.844 |
| 1000 | 10000 | 66600 | 50% | 50% | 50 | 303.5 | 0.996 | 0.977 | 0.966 | 0.824 |
| 1000 | 10000 | 66600 | 65% | 35% | 10 | 63.2 | 0.997 | 0.980 | 0.983 | 0.851 |
| 1000 | 10000 | 66600 | 65% | 35% | 30 | 187.3 | 0.997 | 0.979 | 0.973 | 0.836 |
| 1000 | 10000 | 66600 | 65% | 35% | 50 | 304.2 | 0.995 | 0.973 | 0.963 | 0.809 |
| 1000 | 10000 | 99900 | 35% | 65% | 10 | 98.1 | 0.998 | 0.989 | 0.990 | 0.921 |
| 1000 | 10000 | 99900 | 35% | 65% | 30 | 286.9 | 0.998 | 0.990 | 0.984 | 0.897 |
| 1000 | 10000 | 99900 | 35% | 65% | 50 | 466.2 | 0.996 | 0.981 | 0.977 | 0.863 |

| | | | | | | | | | | |
|---|---|---|---|---|---|---|---|---|---|---|
| 1000 | 10000 | 99900 | 50% | 50% | 10 | 99.0 | 0.998 | 0.988 | 0.993 | 0.926 |
| 1000 | 10000 | 99900 | 50% | 50% | 30 | 287.8 | 0.999 | 0.991 | 0.989 | 0.906 |
| 1000 | 10000 | 99900 | 50% | 50% | 50 | 463.3 | 0.997 | 0.984 | 0.977 | 0.876 |
| 1000 | 10000 | 99900 | 65% | 35% | 10 | 98.9 | 1.000 | 0.995 | 0.992 | 0.928 |
| 1000 | 10000 | 99900 | 65% | 35% | 30 | 287.9 | 0.999 | 0.991 | 0.984 | 0.893 |
| 1000 | 10000 | 99900 | 65% | 35% | 50 | 463.7 | 0.996 | 0.983 | 0.976 | 0.862 |
| 1000 | 15000 | 49950 | 35% | 65% | 10 | 46.4 | 0.998 | 0.977 | 0.968 | 0.822 |
| 1000 | 15000 | 49950 | 35% | 65% | 30 | 136.4 | 0.994 | 0.970 | 0.963 | 0.826 |
| 1000 | 15000 | 49950 | 35% | 65% | 50 | 221.3 | 0.992 | 0.963 | 0.956 | 0.810 |
| 1000 | 15000 | 49950 | 50% | 50% | 10 | 45.9 | 1.000 | 0.981 | 0.963 | 0.845 |
| 1000 | 15000 | 49950 | 50% | 50% | 30 | 136.4 | 0.995 | 0.974 | 0.960 | 0.830 |
| 1000 | 15000 | 49950 | 50% | 50% | 50 | 223.1 | 0.992 | 0.968 | 0.959 | 0.830 |
| 1000 | 15000 | 49950 | 65% | 35% | 10 | 45.8 | 1.000 | 0.977 | 0.964 | 0.838 |
| 1000 | 15000 | 49950 | 65% | 35% | 30 | 137.1 | 0.995 | 0.975 | 0.964 | 0.841 |
| 1000 | 15000 | 49950 | 65% | 35% | 50 | 223.3 | 0.991 | 0.965 | 0.960 | 0.825 |
| 1000 | 15000 | 99900 | 35% | 65% | 10 | 97.2 | 1.000 | 0.999 | 0.994 | 0.931 |
| 1000 | 15000 | 99900 | 35% | 65% | 30 | 289.8 | 1.000 | 0.997 | 0.989 | 0.910 |
| 1000 | 15000 | 99900 | 35% | 65% | 50 | 468.4 | 0.998 | 0.990 | 0.980 | 0.888 |
| 1000 | 15000 | 99900 | 50% | 50% | 10 | 98.0 | 1.000 | 0.998 | 0.988 | 0.935 |
| 1000 | 15000 | 99900 | 50% | 50% | 30 | 287.4 | 1.000 | 0.997 | 0.989 | 0.917 |
| 1000 | 15000 | 99900 | 50% | 50% | 50 | 469.1 | 0.998 | 0.992 | 0.983 | 0.902 |
| 1000 | 15000 | 99900 | 65% | 35% | 10 | 97.3 | 1.000 | 0.998 | 0.993 | 0.940 |
| 1000 | 15000 | 99900 | 65% | 35% | 30 | 288.4 | 0.999 | 0.996 | 0.986 | 0.904 |
| 1000 | 15000 | 99900 | 65% | 35% | 50 | 469.4 | 0.997 | 0.989 | 0.980 | 0.890 |
| 1000 | 15000 | 149850 | 35% | 65% | 10 | 152.2 | 1.000 | 1.000 | 0.993 | 0.946 |
| 1000 | 15000 | 149850 | 35% | 65% | 30 | 444.7 | 1.000 | 0.999 | 0.995 | 0.944 |
| 1000 | 15000 | 149850 | 35% | 65% | 50 | 716.2 | 0.999 | 0.996 | 0.989 | 0.926 |
| 1000 | 15000 | 149850 | 50% | 50% | 10 | 152.0 | 1.000 | 0.999 | 0.998 | 0.959 |
| 1000 | 15000 | 149850 | 50% | 50% | 30 | 444.5 | 1.000 | 0.999 | 0.997 | 0.953 |
| 1000 | 15000 | 149850 | 50% | 50% | 50 | 715.7 | 0.999 | 0.997 | 0.991 | 0.939 |
| 1000 | 15000 | 149850 | 65% | 35% | 10 | 153.0 | 1.000 | 0.999 | 0.993 | 0.958 |
| 1000 | 15000 | 149850 | 65% | 35% | 30 | 445.0 | 1.000 | 0.997 | 0.994 | 0.943 |
| 1000 | 15000 | 149850 | 65% | 35% | 50 | 717.6 | 0.999 | 0.995 | 0.989 | 0.926 |
